# Epigenomic profiling discovers trans-lineage SOX2 partnerships driving tumor heterogeneity in lung squamous cell carcinoma

**DOI:** 10.1101/646034

**Authors:** Takashi Sato, Seungyeul Yoo, Ranran Kong, Maya Fridrikh, Abhilasha Sinha, Prashanth Chandramani-Shivalingappa, Ayushi Patel, Osamu Nagano, Takashi Masuko, Mary Beth Beasley, Charles A. Powell, Jun Zhu, Hideo Watanabe

## Abstract

Molecular characterization of lung squamous cell carcinoma (LUSC), a major subtype of lung cancer, has not sufficiently improved its non-stratified treatment strategies over decades. Accumulating evidence suggests that lineage-specific transcriptional regulators control differentiation states during cancer evolution, and underlie their distinct biological behaviors. In this study, by investigating the super-enhancer landscape of LUSC, we identified a previously undescribed ‘neural’ subtype defined by Sox2 and a neural lineage factor Brn2, as well as the classical LUSC subtype defined by Sox2 and its classical squamous partner p63. Robust protein-protein interaction and genomic co-occupancy of Sox2 and Brn2, in place for p63 in the classical LUSC, indicated their transcriptional cooperation imparting this unique lineage state in the ‘neural’ LUSC. Forced expression of p63 downregulated Brn2 in the ‘neural’ LUSC cells and invoked the classical LUSC lineage with more squamous/epithelial features, which were accompanied by increased activities of ErbB/Akt and MAPK-ERK pathways suggesting differential dependency. Collectively, our data demonstrate heterogeneous cell lineage states of LUSC featured by Sox2 cooperation with Brn2 or p63, for which distinct therapeutic approaches may be warranted.

## Introduction

Lung cancer is the leading cause of cancer-related death worldwide (Bray et al. 2018). Despite recent progress in diagnosis and treatment including molecular-targeted therapeutics and immunotherapy which provide a considerable survival benefit for lung cancer patients, the overall 5-year survival remains less than adequate at 19% (Siegel et al. 2019). Lung squamous cell carcinoma (LUSC) is the second most common histological subtype of lung cancer and is strongly associated with cigarette smoking. Although achievements in tobacco control in developed countries have contributed to a decline in mortality rate of lung cancer, LUSC still remains a major cause of death worldwide and shows even higher incidence than lung adenocarcinoma in several countries (Islami et al. 2015; Cheng et al. 2016). Comprehensive genomic characterization of LUSC conducted by The Cancer Genome Atlas (TCGA) project revealed its heterogeneous features with complex genomic alterations (Cancer Genome Atlas Research Network 2012). Unfortunately, these findings have not resulted in the successful development of clinically approved targeted drugs for LUSC. Compared to patients with lung adenocarcinoma, where multiple genome-based targeted therapeutics have been approved, LUSC patients have very limited therapeutic options. More recently, immune checkpoint inhibitors have emerged as key therapeutic options for solid tumors including LUSC (Brahmer et al. 2015; Herbst et al. 2016); however, the response rate for unselected LUSC population remains only approximately 20% and initially-responded tumors eventually progress in most cases. Therefore, innovative strategies to better characterize and classify LUSC for a better patient stratification for current and future therapeutic options are desperately needed.

We previously identified *SOX2* as the most commonly amplified oncogene in LUSC (Bass et al. 2009). We proposed *SOX2* as a lineage-survival oncogene in squamous cell cancers for its essential role during the development in the specification of the squamous cell lineages by opposing the role of Nkx2-1 in the dividing foregut and its essentiality for LUSC cell survival (Que et al. 2007). In a following study, we identified another squamous lineage factor, p63 as an important cooperative partner of Sox2 in LUSC (Watanabe et al. 2014). *SOX2* amplification on chromosome 3q in LUSC often extends to its telomeric side to include the locus of *TP63*, which encodes p63. While focal amplification and/or overexpression of both *SOX2* and *TP63* genes is found in only 7 % of LUSC tumors, broader copy number gains on 3q telomeric ends are observed in the vast majority of LUSCs (Cancer Genome Atlas Research Network 2012). Studies on expression profiles classified LUSCs into four expression subtypes (primitive, classical, secretory, and basal), suggesting the heterogeneity of transcriptional programs within LUSCs (Wilkerson et al. 2010; Cancer Genome Atlas Research Network 2012; Wu et al. 2013). Based on this classification, co-amplification of *SOX2* and *TP63* is enriched in the classical subtype of LUSC whereas *TP63* expression is relatively low in the primitive and secretory subtypes. However, it remains largely unknown what mechanisms are involved in controlling transcriptional programs in the heterogeneous group of LUSCs.

A series of recent genome-wide histone modification analyses have demonstrated the presence of large clusters of putative enhancers in close genomic proximity, coined super-enhancers (Hnisz et al. 2013; Pott and Lieb 2015). Since these regions typically exhibit cell-lineage-specific patterns in health and in disease, super-enhancer profiling is becoming a powerful tool to identify novel cancer cell lineages governed by specific transcriptional regulators. For example, analysis of the super-enhancer landscape of neuroblastoma has demonstrated heterogeneity of lineage states governed by specific transcriptional programs that may underlie the cause of relapse after chemotherapy (Boeva et al. 2017). Similar analysis of the super-enhancer landscape in acute myeloid leukemia identified a novel epigenomic subtype, which have therapeutic implication for differentiation therapy (McKeown et al. 2017). In the present study, we investigated super-enhancer profiles in LUSC and identified a novel subtype in which Sox2 and a neural transcription factor Brn2 have key roles in determining its distinctive differentiation state. In addition, we show that Brn2 serves as an interacting partner for Sox2 in this novel subtype, instead of p63 in the classical subtype of LUSC and that forced expression of p63 leads to classical squamous cell differentiation and suppression of Brn2 in this novel LUSC subtype.

## Results

### Super-enhancer profiling identifies a novel subtype of lung squamous cell carcinoma

To understand the inter-tumor heterogeneity of cell lineages in LUSC, we examined super-enhancer landscape in a panel of 13 LUSC cell lines. Unsupervised hierarchical clustering of these cell lines using H3K27 acetylation signals over super-enhancer regions near transcriptional regulators identified three subgroups of LUSC (Fig. 1A). Principal component analysis (PCA) using those signals supported this classification (Supplemental Fig. S1A). One of the three subgroups consists of five LUSC cell lines, in which common super-enhancers are observed at genetic loci of 9 transcriptional regulators including *SOX2* and *TP63* loci (Supplemental Fig. S1B; Supplemental Table S1), consistent with our previous findings that these genes play essential roles as lineage oncogenes in typical LUSCs (Bass et al. 2009; Watanabe et al. 2014). This subgroup is enriched in *SOX2*-amplified LUSCs with high expression of both *SOX2* and *TP63* (Fig. 1B). In contrast, LK2 and NCI-H520 cells formed a small subgroup (Fig. 1A; Supplemental Fig. S1A), in which the common super-enhancers with highest signals lie on only two loci at the *SOX2* gene and the *POU3F2* gene, which encodes Brn2 (Fig. 1C,D; Supplemental Table S2). Brn2 is a well-known lineage factor in neural progenitor cells, particularly in the hypothalamus, where it partners with Sox2 to exert its transcriptional functions (Schonemann et al. 1995; Tanaka et al. 2004; Lodato et al. 2013). We also noted that the third subgroup which consists of 5 cell lines shows relatively low expression of both *SOX2* and *TP63* (Fig. 1B) although human LUSC tumors with low expression (< mean−SD) of both genes accounts for only 6.6 % in the Cancer Genome Atlas Lung Squamous Cell Carcinoma (TCGA-LUSC) dataset. While the *SOX2* super-enhancers were shared by the former two subgroups (Fig. 1C,D; Supplemental Fig. S1B) and the *TP63* super-enhancers were found only in the first subgroup (Fig. 1D; Supplemental Fig. S1B), *POU3F2* super-enhancers were present only in the small subset of LK2 and NCI-H520 (Fig. 1D). We also found the *POU3F2* super-enhancers to be among the top differential super-enhancers in this subset compared to the ‘classical’ SOX2/TP63 subgroup (Supplemental Fig. S1C). Analyses using RNA-seq data on these cell lines obtained through Cancer Cell Line Encyclopedia (CCLE) dataset revealed that expression of *POU3F2* was indeed significantly higher in this small subset while the squamous lineage factor *TP63* was expressed significantly higher in the ‘classical’ *SOX2*/*TP63* subgroup of LUSC (Fig. 1B). By contrast, expression level of *SOX2* was found in both subgroups and highest in the *SOX2*/*POU3F2* subset (Fig. 1B). Immunoblotting showed the expression pattern to be consistent with protein level; the LUSC cell lines that harbor *POU3F2* super-enhancers do not express p63 at a detectable level but have high expression of Sox2 and Brn2 (Fig. 1E). These findings suggest that, while *SOX2* is highly expressed and likely serves as a lineage factor for both of these two subgroups, this small subgroup represents a unique subset of LUSCs signified by the neural lineage factor Brn2.

**Figure 1.**
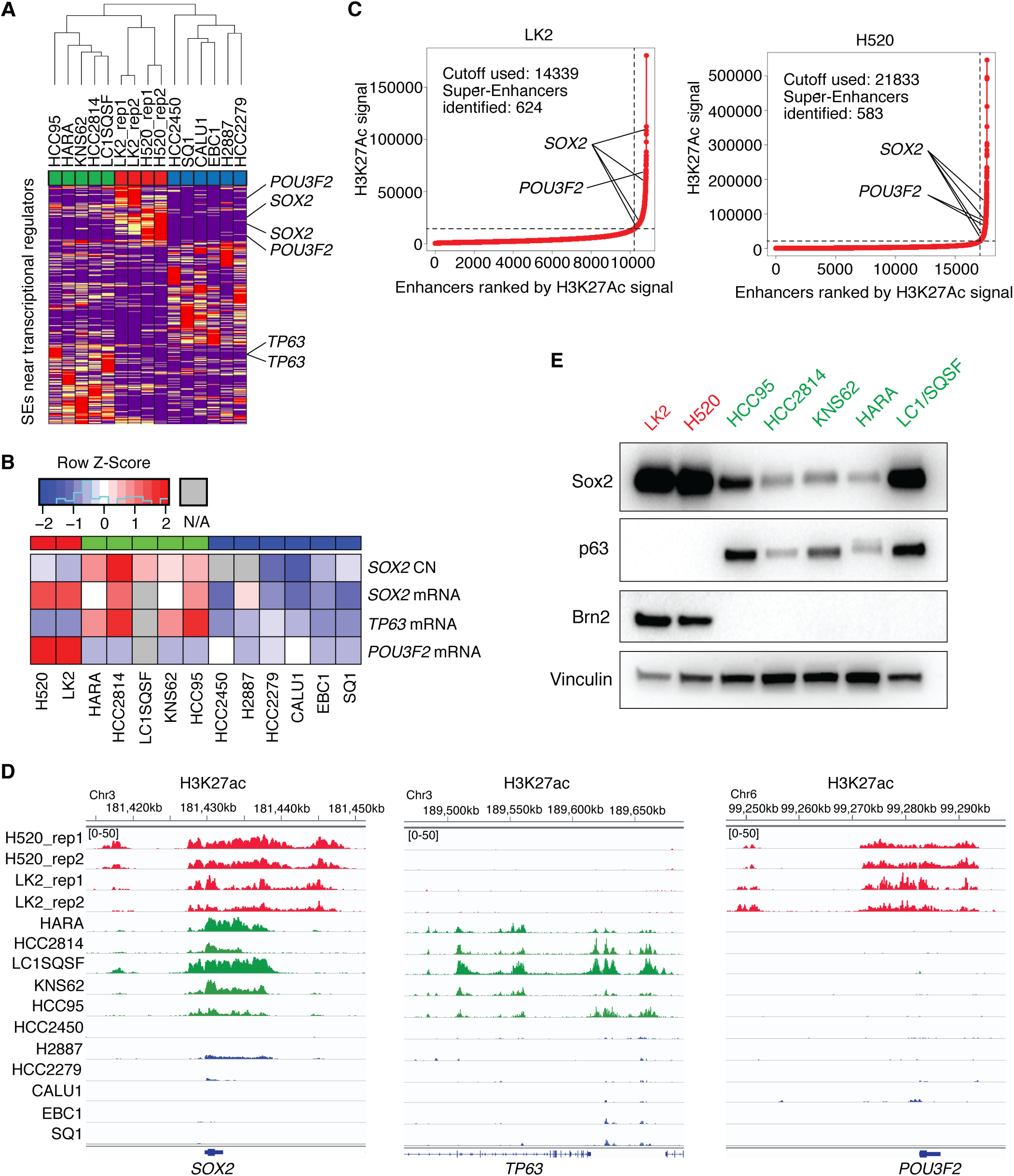
Super-enhancer profiling identifies a novel subtype of LUSC. *(A)* Unsupervised hierarchical clustering of 13 LUSC cell lines using super-enhancer scores near transcriptional regulator genes. *(B)* Copy number of *SOX2* and mRNA expression of *SOX2*, *TP63* and *POU3F2* in LUSC cell lines from CCLE. *(C)* Super-enhancer plots using H3K27ac scores in the small subset of LUSC cell lines; LK2 cells (*left*) and NCI-H520 cells (*right*). *(D)* Genome view tracks of H3K27ac signal at loci of *SOX2* (*left*), *TP63* (*middle*) and *POU3F2* (*right*) in LUSC cell lines. *(E)* Protein expression of Brn2, p63, Sox2 and vinculin as a loading control in LUSC cell lines.

### BRN2 signifies the ‘neural’ subtype in human LUSCs

Brn2 has been described to have an oncogenic role in neuroendocrine tumors such as small cell lung cancer (SCLC), neuroendocrine prostate cancer and glioblastoma (Schreiber et al. 1990; Ishii et al. 2013; Ishii et al. 2014; Suva et al. 2014; Bishop et al. 2017). To determine whether this unique ‘neural’ subset exists in human LUSC data and *POU3F2 (Brn2)* signifies this subgroup, we examined TCGA-LUSC dataset and identified a subset of LUSC that expresses *POU3F2* (Fig. 2A) and their expression levels were comparable to that in SCLC (George et al. 2015) and hypothalamus in GTEx data (GTEx Consortium 2013) (Fig. 2B), suggesting functional relevancy of Brn2 in this unique subset of LUSC. Of note, this ‘neural’ subset of LUSC forms a distinct cluster from super-enhancer profiles of SCLC cell lines (data not shown). In order to investigate how these tumors with Brn2 expression are represented in clinical specimens, we examined Brn2 expression pattern at protein level in human LUSC tissues by immunohistochemical staining. While all 10 specimens were positive for Sox2 as expected, we found Brn2 expression in 2 specimens (Fig. 2C; Supplemental Fig. S2A). Of note, the Brn2 expression was relatively confined to subpopulations of tumor cells in these two LUSC tissues. This suggests that some of the tumors with intermediate *POU3F2* expression level observed in bulk mRNA in the TCGA-LUSC dataset may reflect heterogeneous tumor populations with partial ‘neural’ differentiation and that, perhaps unlike cell lines, distinct lineage states can coexist within the same LUSC tumor. This further implies trans-differentiation during the evolution of LUSCs. We next examined the correlation of expression of *POU3F2* with *SOX2* and *TP63* expression in the TCGA-LUSC dataset. We found that expression of *POU3F2* was anti-correlated with that of *TP63* (Fig. 2B; Supplemental Fig. S2B), consistent with the data from LUSC cell lines in which high expression of *POU3F2* and *TP63* were mutually exclusive. We did not find significant correlation between *POU3F2* and *SOX2* expression (Supplemental Fig. S2C) in the entire dataset; however, when we focused on tumors with low *TP63* expression, we found a significant correlation of *POU3F2* with *SOX2* (Fig. 2D) whereas there was a significant correlation of *TP63* with *SOX2* among LUSCs with high *TP63* expression (Supplemental Fig. S2D). These findings suggest that Brn2 and p63 have contrasting roles representing distinct lineages within LUSC by partnering with Sox2 to elicit distinct transcriptional outputs.

**Figure 2.**
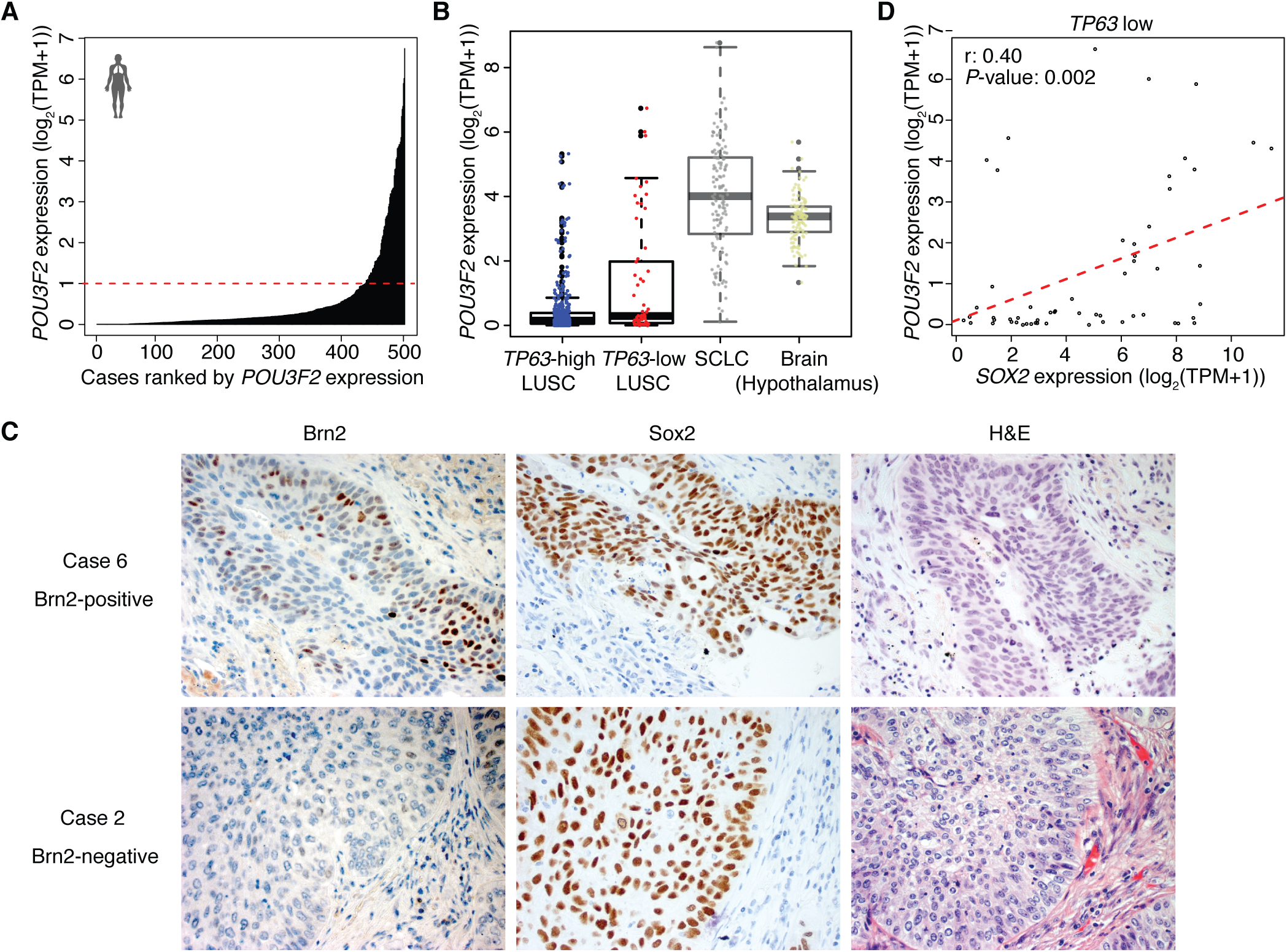
Brn2 is expressed in a subset of human primary LUSC tumors. *(A)* Expression of *POU3F2* in 501 TCGA LUSC tumor tissues. The red dashed line (TPM=1) shows the cutoff to separate samples into *POU3F2* high and low samples. *(B)* Box plots of *POU3F2* expression in *TP63*-high and low LUSC tumors from TCGA, SCLC tumors from publicly available datasets and normal brain tissues (hypothalamus) from GTEx. *(C)* Immunohistochemical staining of Brn2 and Sox2 and H&E staining in Brn2-positive and negative human LUSC tumors. Representative images are shown (original images, ×400).

To investigate the difference in gene signatures between *POU3F2*-expressing LUSC tumors and *TP63*-expressing LUSC tumors, we next performed differential expression analysis in the TCGA-LUSC dataset comparing *POU3F2*-high/ *TP63*-low tumors vs. *TP63*-high/ *POU3F2*-low tumors (Fig. 3A). A functional enrichment analysis of differentially expressed genes identified genes which play roles in neural cell differentiation significantly enriched in *POU3F2*-high/ *TP63*-low tumors while those involved in epithelial/ epidermal cell differentiation were significantly enriched in *TP63*-high/ *POU3F2*-low tumors (Fig. 3B). Further, we evaluated the prognostic significance of *TP63*/ *POU3F2* expression status in the TCGA-LUSC dataset and found that patients with *POU3F2*-high/ *TP63*-low tumors showed significantly shorter survival compared to those with *TP63*-high/ *POU3F2*-low tumors, suggesting clinically distinct, aggressive features of this ‘neural’ subtype (Fig. 3C). These findings indicate the presence of a distinct LUSC subset with neural differentiation that has not been described previously.

**Figure 3.**
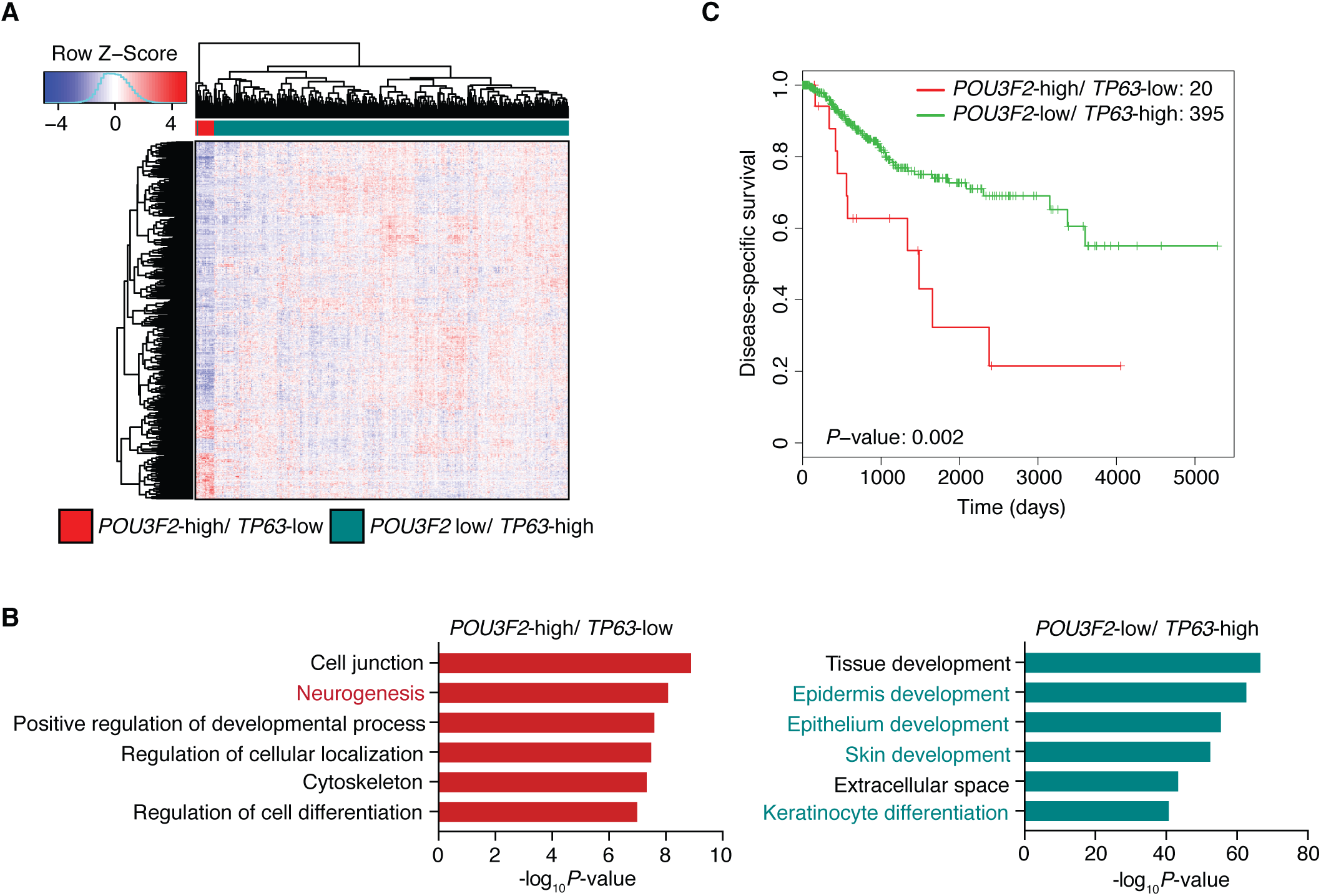
Brn2 signifies the ‘neural’ subtype of LUSC in human primary LUSC tumors. *(A)* Heatmap showing hierarchical clustering of LUSC tumors from TCGA using differentially expressed genes between *POU3F2*-high/ *TP63*-low (n=20) and *POU3F2*-low/ *TP63*-high tumors (n=396). With cutoffs of fold change>2 and FDR<0.01, 196 genes are differentially up-regulated and 735 genes are differentially down-regulated in the *POU3F2*-high/ *TP63*-low tumors. Color scheme represents Z-score distribution. *(B)* Gene ontology analyses for the differentially up-regulated (*left*) and down-regulated (*right*) genes. Enriched functions for these genes are identified based on Fisher’s exact test against GO terms curated in MSigDB. *(C)* Kaplan-Meier curves plotted to compare the disease-specific survival of *POU3F2*-high/ *TP63*-low (n=20) versus *POU3F2*-low/ *TP63*-high (n=395) LUSC tumors from TCGA. Log-rank test *P*-value=0.002.

### BRN2 and SOX2 interact and co-localize at genetic loci in the ‘neural’ LUSC cells

Given the positive correlation of *POU3F2* and *SOX2* expressions in LUSC tumors with low *TP63* expression, and previously reported interaction of Brn2 and Sox2 in neural progenitor cells (Tanaka et al. 2004; Lodato et al. 2013), we further investigated the relationship between Brn2 and Sox2 in LUSC. First, using immunofluorescence analysis we found that Sox2 and Brn2 co-localize in the nuclei of the Brn2-positive LUSC cell lines (Fig. 4A; Supplemental Fig. S3A). In contrast, Sox2 but not Brn2 was detected in nuclei of the ‘classical’ Sox2/p63-positive LUSC cells confirming the specificity of the antibody (Supplemental Fig. S3B). Second, we observed their robust protein-protein interaction by endogenous and reciprocal co-immunoprecipitations in the two ‘neural’ LUSC cell lines (Fig. 4B). Given their physical interaction, we hypothesized that Brn2 and Sox2 co-occupy genomic loci to coordinately regulate gene expression in these Brn2-positive LUSC cells. To test for genomic co-occupancy of Brn2 and Sox2, we explored genome-wide binding profiles of Brn2 and Sox2 in the ‘neural’ LUSC cells by chromatin immunoprecipitation sequencing (ChIP-seq) and found that the Brn2 binding peaks substantially overlap with the peaks for Sox2 in these cells (Fig. 4C,D). In contrast, the Sox2 binding peaks from ‘classical’ LUSC cells showed distinct profiles with less overlaps with those for Brn2 in ‘neural’ LUSC cells (Fig. 4C,D), suggesting that in the absence of p63, Brn2 engages Sox2 to localize at genetic loci thereby imparting ‘neural’ features on LUSC cells.

**Figure 4.**
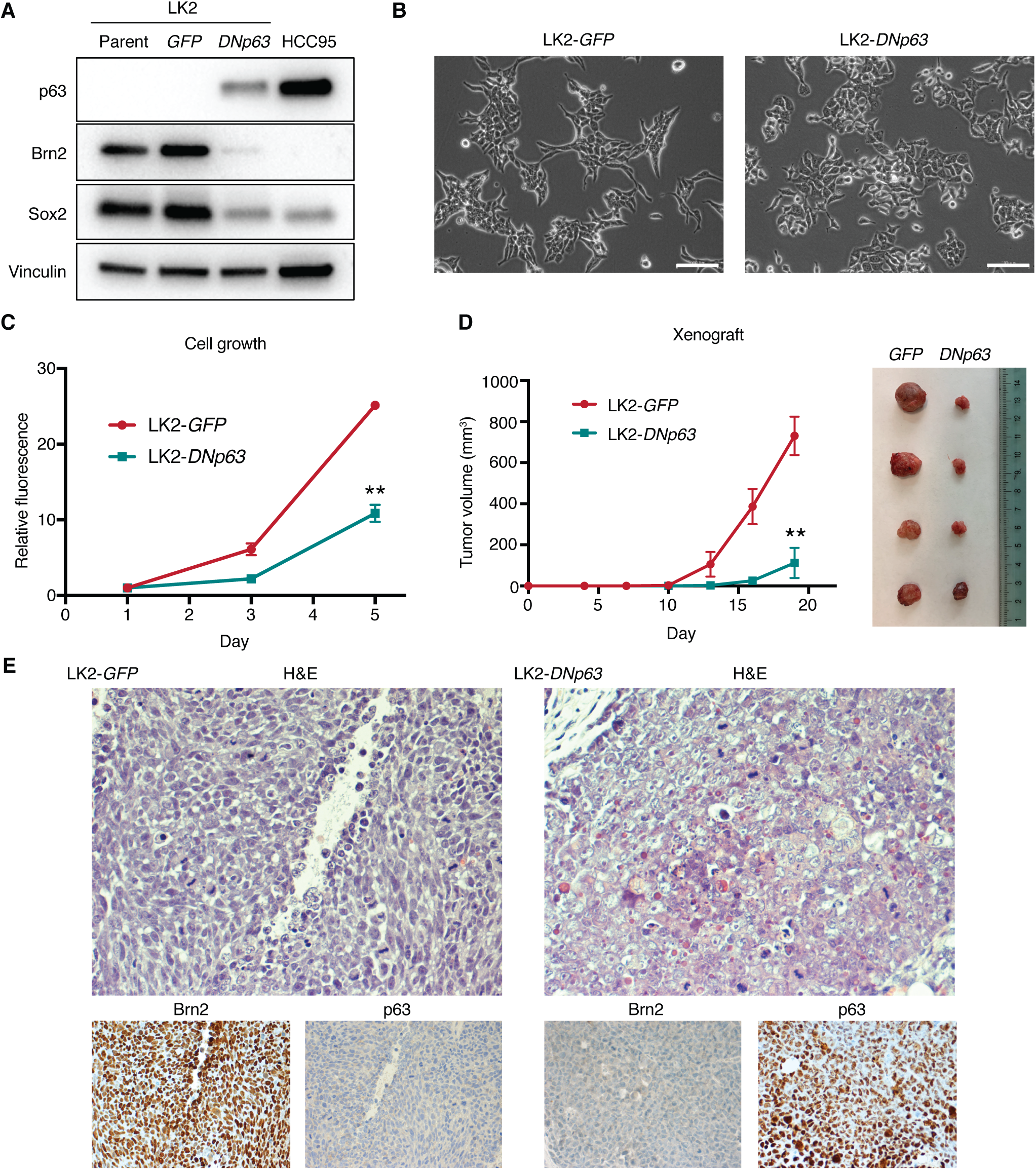
DNp63 overexpression in the ‘neural’ LUSC cells suppresses Brn2 expression and induces phenotypic changes. *(A)* Protein expression of p63, Brn2, Sox2 and vinculin as a loading control in parental, GFP-overexpressed and DNp63-overex-pressed LK2 cells. *(B)* Phase-contrast microphotographs of GFP-overexpressed and DNp63-overexpressed LK2 cells. Bar = 100 μm. *(C)* Cell growth of GFP-overexpressed and DNp63-overexpressed LK2 cells. Mean ± SD of sextuplicates are shown. **, *P*<0.001 vs. GFP-overexpressed LK2 cells, *t*-test. *(D)* Tumor growth of GFP-overexpressed and DNp63-overexpressed LK2 cells *in vivo*. Mean ± SD of tetraplicates are shown. **, *P*<0.001 vs. GFP-overexpressed xenografts, *t*-test. Xenograft tumors resected 19 days after inoculation are shown in the right picture. *(E)* H&E staining and immunohistochemical staining of Brn2 and p63 in GFP-overexpressed or DNp63-overexpressed LK2 xenograft. Original Images, ×400.

### p63 suppresses Brn2 expression and induces a classical-squamous tumor state

We previously reported that p63 and Sox2 physically interact and exhibit overlapping genomic occupancy, which suggest their cooperative transcriptional programs in ‘classical’ LUSCs (Watanabe et al. 2014). The interaction of Brn2, in place of p63, with Sox2 that we observed in this ‘neural’ subset implies counteractive functions of Brn2 and p63 in lineage determination. To date, however, the role of the balance between these two factors in differentiation of squamous tissue, or in any lineages, has not been described. Therefore, to determine whether p63 has any effects on Brn2 or vice versa, we first overexpressed deltaNp63-alpha (DNp63), the predominant isoform in squamous lineage and LUSC, in the ‘neural’ LUSC cells (Fig. 5A; Supplemental Fig. S4A,B). Introduction of DNp63 substantially suppressed Brn2 expression, while it also suppressed expression of Sox2 to a lesser extent in these cells (Fig. 5A; Supplemental Fig. S4A,B). Notably, DNp63 overexpression led to morphological changes in these ‘neural’ LUSC cells from small and/or elongated morphology to larger and cuboidal morphology in conventional 2-dimensional culture (Fig. 5B; Supplemental Fig. 5C) and decreased cell proliferation (Fig. 5C; Supplemental Fig. S4D). To confirm whether these observed changes *in vitro* reflect a shift in differentiation status, we next investigated the effects of DNp63 over-expression on the ‘neural’ LUSC *in vivo*. In the xenograft model, the tumor growth rates were significantly decreased in DNp63-overexpressed LUSC xenografts compared to control ‘neural’ LUSC (Fig. 5D). In addition, Brn2 expression was suppressed in p63-positive tumor cells (Fig. 5D; Supplemental Fig. S4E,F). Consistent with our finding *in vitro*, DNp63 expression downregulated expression of Sox2 (Supplemental Fig. S4D,F,G) presumably reflecting the squamous epithelial differentiation induced by p63 (Truong and Khavari 2007; Arnold et al. 2011). Notably, patches of p63-negative tumor cells in DNp63-overexpressed xenograft tumor shows reduced expression of Brn2 and Sox2 (Supplemental Fig. S4E,F), likely due to imperfect selection or heterogeneous ectopic expression of DNp63, exhibiting a clear effect of DNp63 on suppressing Brn2. Morphologically, p63-positive tumor cells showed open chromatin, larger nucleoli, and on average, had a polygonal shape and more cytoplasm indicative of more classical squamous cell carcinoma histology while p63-negative/ Brn2-positive cells have a slightly oval shape, less cytoplasm, denser chromatin and less frequent/smaller nucleoli (Fig. 5D; Supplemental Fig. S4F). These results suggest that DNp63 counteracts with Brn2 and induces a different tumor state characterized by more squamous/epithelial features of the cells.

**Figure 5.**
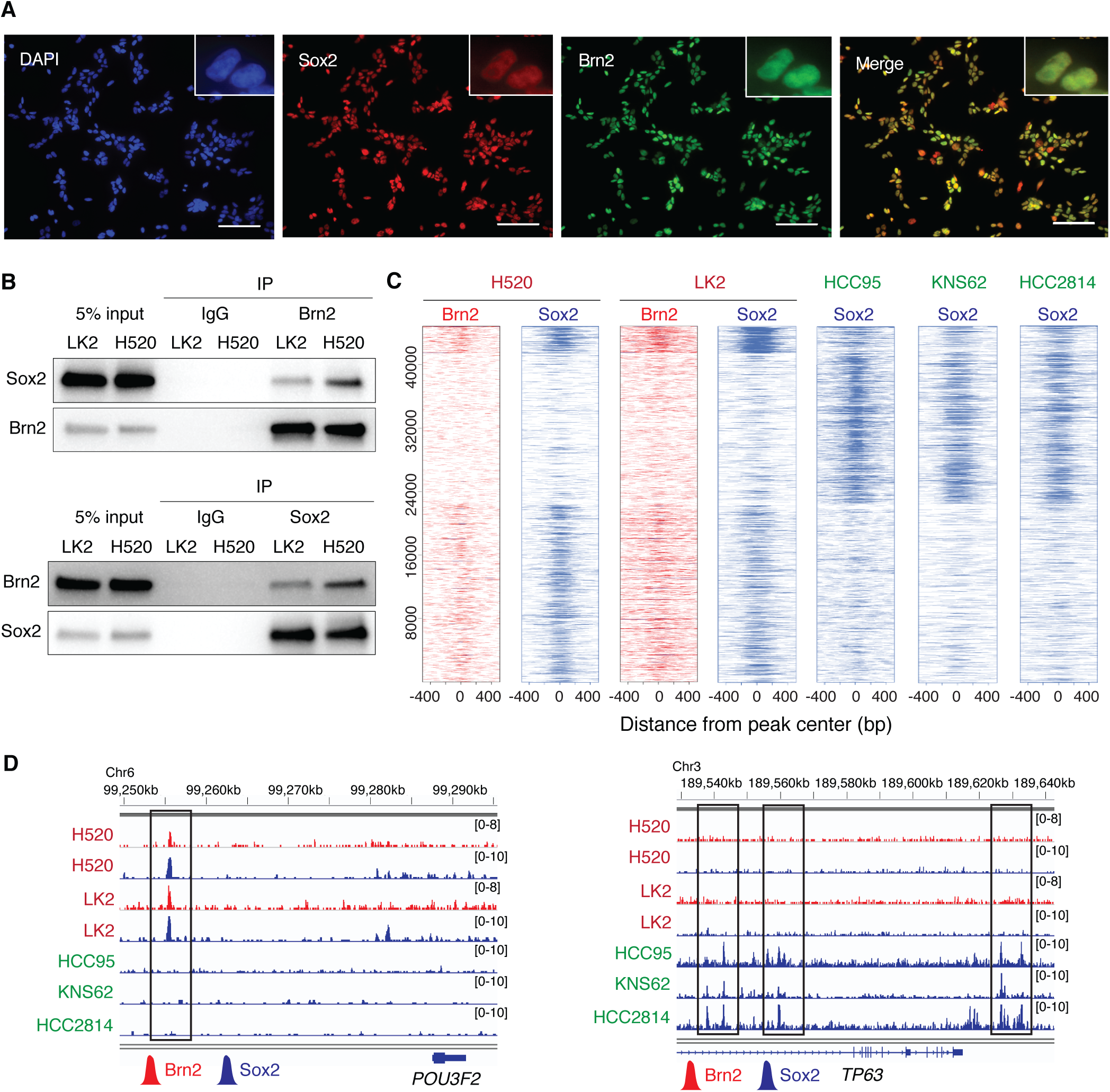
Brn2 and Sox2 interact and co-localize at genetic loci in the ‘neural’ subset of LUSC. *(A)* Expression of endogenous Brn2 and Sox2 in LK2 cells, determined by immunofluorescence with anti-Sox2 (*green*) and anti-Brn2 (*red*) antibodies, respectively. DAPI staining (nuclei; *blue*) and merged images are also shown. Original magnification, ×200. Scale bar, 100 μm. *(B)* Sox2-Brn2 interaction, shown by co-immunoprecipitation of Sox2 using an antibody against endogenous Brn2 (*top*) and co-immunoprecipitation of Brn2 using an antibody against endogenous Sox2 (*bottom*) in LK2 and NCI-H520 cells. *(C)* Heatmap depicting global analysis of ChIP-seq signals for Brn2 and Sox2 in ‘neural’ NCI-H520 and LK2 cells and those for Sox2 in ‘classical’ HCC95, KNS62 and HCC2814 cells at all the peak loci. ChIP-seq signal intensity is shown by color shading. *(D)* Genome view tracks of Brn2 and Sox2 ChIP-seq signals in NCI-H520 and LK2 cells and Sox2 ChIP signals in HCC95, KNS62 and HCC2814 cells at loci of *POU3F2* (*left*) and *TP63* (*right*).

On the other hand, overexpression of Brn2 in p63-positive ‘classical’ LUSC cells did not lead to suppression of p63 while modestly increased expression of Sox2, indicating non-reciprocal regulatory roles of Brn2 and DNp63, and regulation of Sox2 by Brn2 (Supplemental Fig. S5A,B). To further investigate the role of Brn2 in the ‘neural’ LUSC cells, we ablated expression of Brn2 in these cells by CRISPR-Cas9 mediated deletion of the *POU3F2* gene (Supplemental Fig. S5C). Brn2 ablation led to decreased cell proliferation in ‘neural’ LK2 cells (Supplemental Fig. S5D), suggesting its role in maintaining the lineage state; however, it did not lead to changes in p63 expression (Supplemental Fig. 5C), again suggesting that p63 expression is not under the regulation of Brn2. Although Brn2 overexpression led to increased Sox2 expression, Brn2 ablation did not result in suppression of Sox2 (Supplemental Fig. S5C). Instead, p63 ablation increased expression of Sox2 (Supplemental Fig. S5E), suggesting negative regulation of Sox2 by p63, consistent with the findings that p63-high LUSC cells have lower expression of Sox2 than the Brn2-positive subset (Fig. 1E) and DNp63 induction suppressed SOX2 expression in the same subset (Fig. 5A; Supplemental Fig. S4A,B).

### DNp63 changes the transcriptional program of the ‘neural’ LUSC cells

To investigate how DNp63 changed transcriptional programs leading to the phenotypical changes in ‘neural’ LK2 cells, we performed transcriptomic profiling on these engineered cells (Fig. 6A). We found that genes which play roles in neuronal functions were enriched in down-regulated genes while genes involved in epithelial/ epidermal cell development were enriched in up-regulated genes (Fig. 6A). When we considered the levels of DNp63 expression depending on the experimental models, in addition to neuronal genes, those involved in cell cycle/ DNA replication were also enriched in down-regulated genes (Supplemental Fig. S6A). Consistent with this finding, higher proportion of cells in the S phase of cell cycle was found in control LK2 cells compared to the DNp63-overexpressed cells (Supplemental Fig. S6B,C). Altered cell-cycle dynamics might be a simply reflection of faster cell proliferation but it is also plausible that it is associated with neuronal differentiation in these ‘neural’ LUSC cells, which needs further investigation (Ferguson and Slack 2001; Gobert et al. 2009; Magri et al. 2014). Of note, we found strong associations (OR=9.81 and 6.17 for up- and down-regulated genes, respectively) between gene signatures in this model and those from human TCGA-LUSC tumors based on *POU3F2*/ *TP63* expression (Fig. 3A; Supplemental Table S3), suggesting biological relevancy of our model in the context of human LUSCs. To see whether introduction of DNp63 reconstitutes Sox2-binding profile toward classical LUSC, we profiled genomic occupancy of Sox2 in the DNp63-overexpressed cells and compared to the control ‘neural’ LK2 cells. We found that while average ChIP-seq signals of all the Sox2 peaks decreased in the DNp63-overexpressed cells compared to the control LK2 cells, the DNp63-overexpressed cells showed increased signals near the Sox2 peaks exclusive to the ‘classical’ LUSC lines (Fig. 6C), suggesting that presence of DNp63 re-engages Sox2 cistrome in the ‘classical’ LUSC cells specific Sox2-binding regions (Fig. 6D). Furthermore, we found p63 binding profile formed in the DNp63-overexpressed cells similar to that in ‘classical’ HCC95 cells including overlaps with Sox2 binding peaks from these cells (Fig. 6D; Supplemental Fig. S6D). These findings suggest that DNp63 overrides Sox2’s transcriptional programs in the ‘neural’ state to redefine it into ‘classical’ squamous-cell state via superseding Brn2 as a Sox2 partner.

**Figure 6.**
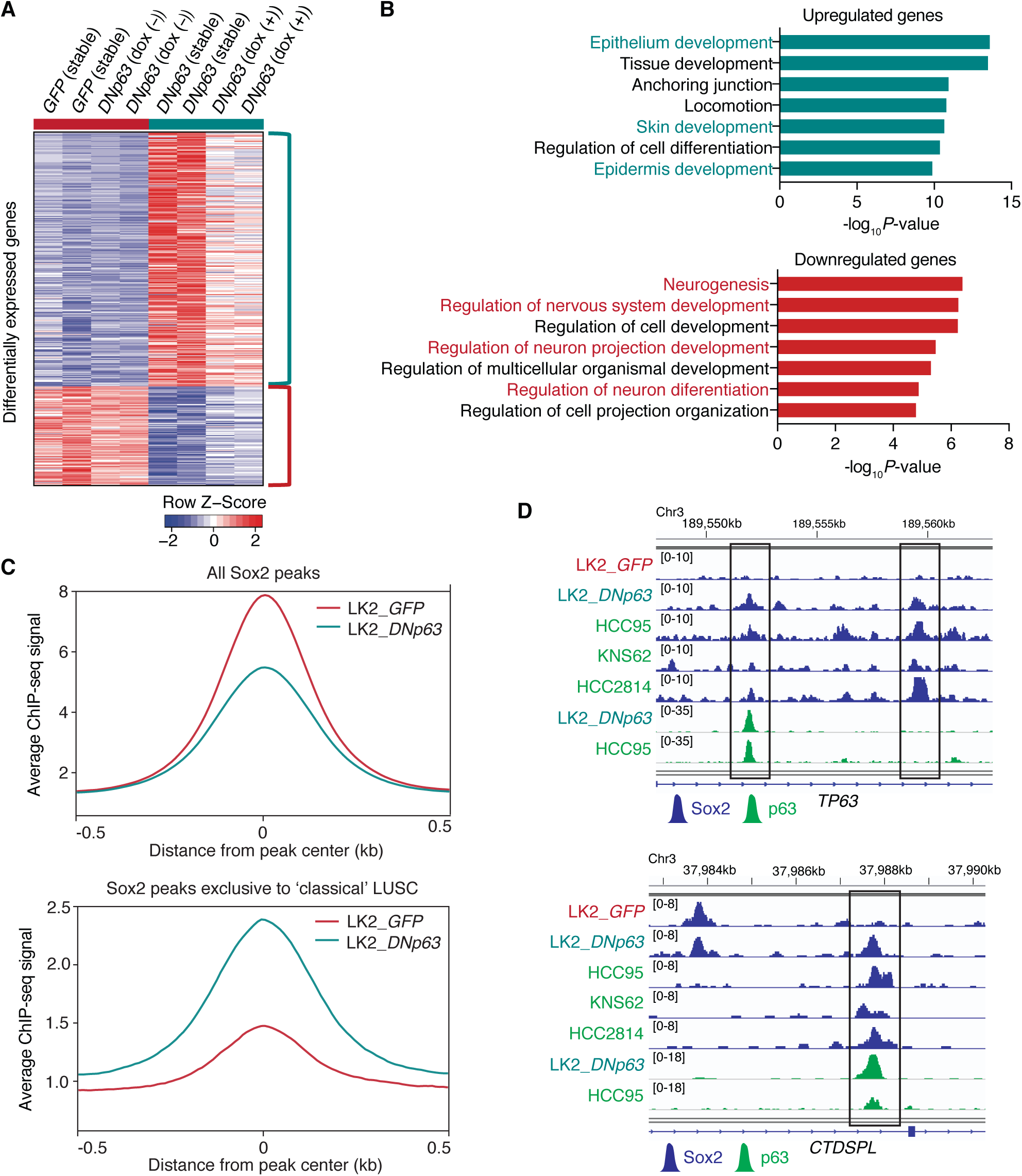
DNp63 induces a classical squamous-cell transcriptional program in the ‘neural’ LUSC cells. *(A)* Heatmap showing 399 differentially expressed genes (287 upregulated and 112 downregulated genes) between control LK2 cells (stable GFP-overexpressed cells and doxycycline-inducible DNp63-overexpressing cells without doxycycline in duplicate) and DNp63-overexpressed LK2 cells (stable DNp63-overexpressed cells and doxycycline-inducible DNp63-overexpressing cells with 2 μg/ml doxycycline in duplicate), sorted by fold change. Color scheme represents Z-score distribution. *(B)* Gene ontology analyses for the differentially up-regulated (*top*) and down-regulated (*bottom*) genes. Enriched functions for these genes are identified based on Fisher’s exact test against GO terms curated in MSigDB. *(C)* Average ChIP-seq signals for all Sox2 peaks (*top*) and Sox2 peaks exclusively found in ‘classical’ HCC95, KNS62 and HCC2814 cells (*bottom*) in GFP-overexpressed and DNp63-overexpressed LK2 cells. *(D)* Genome view tracks of Sox2 ChIP-seq signals in GFP-overexpressed and DNp63-overexpressed LK2 cells, HCC95, KNS62 and HCC2814 cells and p63 ChIP signals in DNp63-overexpressed LK2 cells and HCC95 cells at loci of *TP63* (*top*) and *CTDSPL* (*bottom*).

### DNp63 changes activities of ErbB family signaling in the ‘neural’ LUSC cells

Distinct patterns of cell signaling landscape including receptor tyrosine kinase (RTK) activity are associated with cell differentiation states. To investigate whether DNp63-overexpression induces a differential signaling activity profile in the ‘neural’ LUSC cells, we assayed for human phospho-RTK array on these cells (Fig. 7A). At the original state in the control cells, we detected ErbB4 phosphorylation along with activation of EGFR and IGF1R. In contrast, ErbB4 phosphorylation was diminished and instead higher signals of ErbB3 phosphorylation emerged in the DNp63-overexpressed cells.

**Figure 7.**
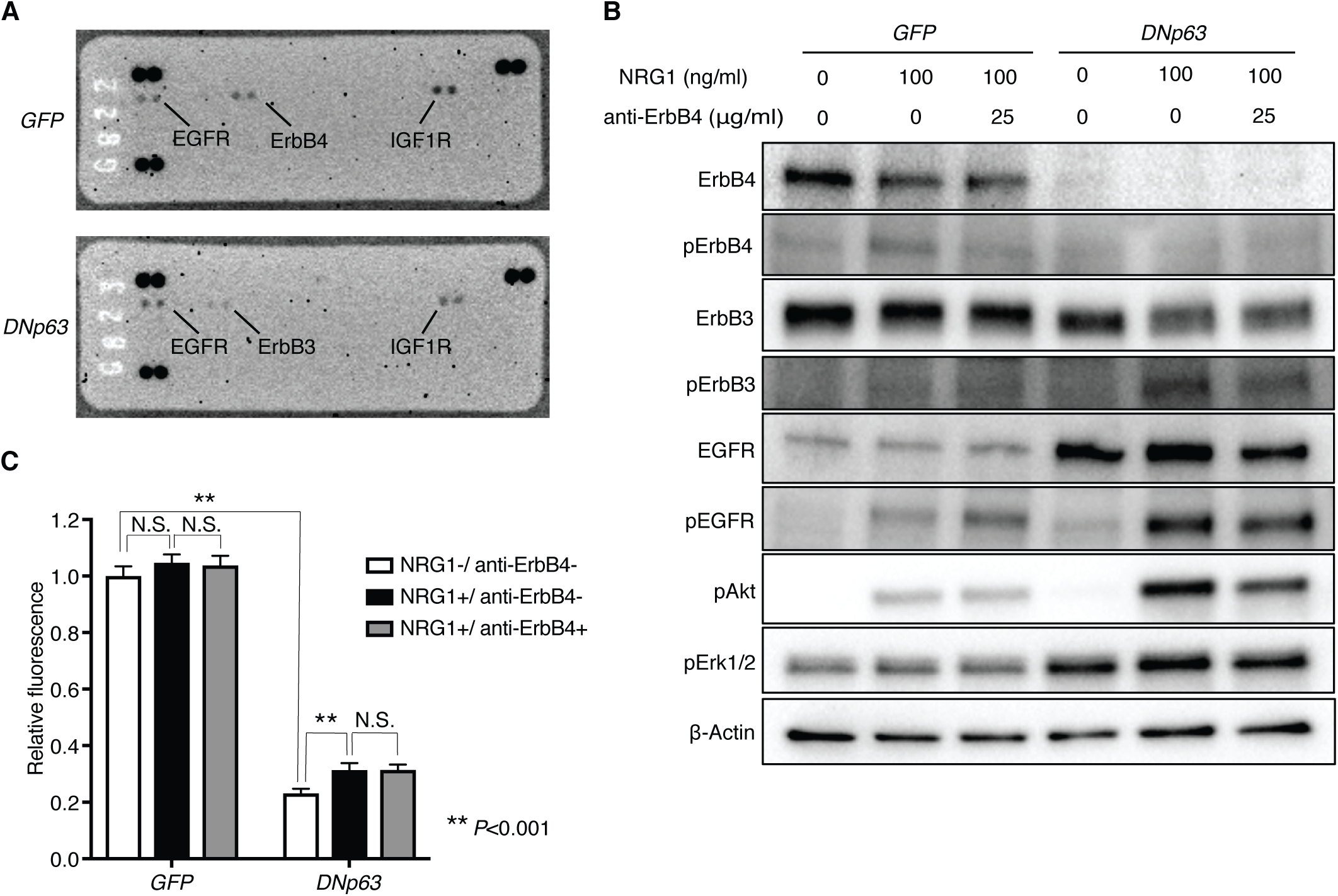
DNp63 alters ErbB family signaling profile in the ‘neural’ LK2 cells. *(A)* Human phospho-receptor tyrosine kinase array was performed for GFP-overexpressed and DNp63-overexpressed LK2 cells. *(B)* Protein expression of ErbB4, phospho-ErbB4, ErbB3, phospho-rbB3, EGFR, phospho-EGFR, phospho-Akt and phospho-Erk1/2 and β-Actin as a loading control in GFP-overexpressed and DNp63-overexpressed LK2 cells. Cells were serum-starved for 4 hours, and then incubated with anti-ErbB4 monoclonal antibody at the indicated concentrations for 15 minutes, followed by stimulation with NRG1 for 30 minutes at the indicated concentrations. *(C)* Cell proliferation of GFP-overexpressed and DNp63-overexpressed LK2 cells. Cells were cultured with 5% FBS and with or without NRG-1 (100 ng/ml) and anti-ErbB4 monoclonal antibody (25 μg/ml) for 4 days. Mean ± SD of sextuplicates are shown. **, *P*<0.001 and N.S., not significant (*P*>0.05), *t*-test with Bonferroni correction..

We confirmed this switch in activity between ErbB family members by immunoblotting and further examined the difference in ErbB family signaling with the use of recombinant Neuregulin-1 (NRG1), a ligand for ErbB3 and ErbB4, as well as a neutralizing antibody directed against ErbB4 (Fig. 7B; Supplemental Fig. 7A). This line of experiments revealed that diminished phosphorylation of ErbB4 upon DNp63 overexpression was a consequence of suppressed ErbB4 protein level. In contrast, expression levels of ErbB3 are similar between DNp63 overexpressed and control cell lines while its activity was significantly increased upon DNp63 overexpression under the same growth condition (10% FBS) (Supplemental Fig. 7A), suggesting that DNp63 expression led to altered expression levels of its ligands or expression/activity of its heterodimeric partners and/or interacting signaling molecules. Indeed, we observed increased protein level of EGFR upon DNp63 overexpression, and in response to NRG1 stimulation, phosphorylation of ErbB3 and EGFR increased to even higher level in the DNp63-overexpressed cells (Fig. 7B; Supplemental Fig. 7A), suggesting that increased EGFR protein level and their homo/hetero-dimerization led to increased phosphorylation of ErbB3 and EGFR. We also found that p63 binds to the *EGFR* locus in the DNp63 over-expressed cells suggesting a direct regulation of *EGFR* gene by p63 at the transcriptional level (Supplemental Fig. S7B).

To determine which major downstream signal pathway mediates ErbB3/EGFR activation, we examined phosphorylation levels of Akt and Erk1/2 in these cells. We observed increased Akt phosphorylation levels corresponding to phosphorylation levels of ErbB3 and EGFR in response to NRG1 stimulation (Fig. 7B; and Supplemental Fig. S7A). In contrast, Erk1/2 phosphorylation levels were higher in the DNp63-overexpressed cells, but did not change after NRG1 stimulation, suggesting that the activation of MAPK-ERK pathway by DNp63 overexpression was independent from the phosphorylation of ErbB3 and EGFR. We next examined response to NRG1 stimulation on cell growth in these cells and found significantly higher response in the DNp63-overexpressed cells compared to the control cells (Fig. 7C). These data suggest that the ‘neural’ LUSC cells are less dependent on ErbB/Akt and MAPK-ERK pathways compared to the more epithelial cell differentiation state induced by DNp63, and further imply different therapeutic strategies could be applied by targeting different molecules depending on the cell differentiation status.

## Discussion

In this report, we identified a novel subtype of LUSC characterized by the neural transcription factor Brn2. We examined super-enhancer landscape of LUSC cell lines to reveal the heterogeneity of transcriptional programs in LUSC and identified three subgroups of LUSC based on the super-enhancer profiles. Notably, super-enhancers on the locus of the *SOX2* gene are shared by two subgroups. We previously reported that *SOX2* is the most commonly amplified gene in LUSC and its expression is essential for cell survival (Bass et al. 2009). Following our study, it has been confirmed that Sox2 promotes lung squamous cancer lineage in mice model (Ferone et al. 2016; Tata et al. 2018), supporting its important role as a lineage oncogene. However, in one subgroup of LUSC, *SOX2* super-enhancers were not found and its expression was low, suggesting that Sox2 does not necessarily govern transcriptional programs among all LUSCs. While this subset is not represented as a significant proportion of primary human LUSC in TCGA cohort, it remains to be seen which lineage state this low Sox2 LUSC subset represents and which transcriptional program may be active. In addition, super-enhancers on the locus of the *TP63* gene, which we also identified as a collaborating lineage factor with Sox2 in LUSC, were only found in one ‘classical’ subgroup of LUSC, suggesting that in the other subgroups, other factors play roles in controlling their lineage states. In fact, our super-enhancer profiling found that super-enhancers on the loci of *SOX2* and *POU3F2* instead of *TP63* were commonly shared in a small subset of LUSC. Brn2, encoded by *POU3F2*, is a Class III POU transcription factor which play an essential role in neural cell differentiation (Sugitani et al. 2002; Jin et al. 2009; Iwafuchi-Doi et al. 2011; Dominguez et al. 2013) and is highly expressed in glioblastoma, neuroendocrine SCLC and neuroendocrine prostate cancer (Schreiber et al. 1990; Ishii et al. 2014; Bishop et al. 2017). This subset exhibited gene expression patterns associated with neural development compared to the ‘classical’ subtype, suggesting the differentiation states of the subset is closer to neural lineages. This subset is not categorized as a neuroendocrine lung cancer which also commonly expresses Brn2 and is close to neural lineages. We speculate that this unique LUSC subset can have a potential to show neuroendocrine phenotypes possibly with additional factors implicated for neuroendocrine lineages such as Rb1 and Ascl1, which warrants further investigation.

In this study, we highlighted the partnership of Sox2 and Brn2 in this unique ‘neural’ subset of LUSC for regulating their transcriptional programs. Sox2 has been described to have a role in determining cell differentiation states by cooperating with other lineage factors such as Oct4 in embryonic stem cells (Boyer et al. 2005), Pax6 in eye lens development (Kamachi et al. 2001), p63 in squamous lineages (Watanabe et al. 2014) and Brn2 in neural progenitor cells (Tanaka et al. 2004; Lodato et al. 2013). Our data support distinct contributions of Sox2 across multiple cell types and confirmed that cooperation of Sox2 and Brn2 in neural progenitor cells is conserved in the ‘neural’ LUSC cells, which has not been described previously.

Prior studies including ours have reported a role of p63 as a lineage oncogene in defining squamous lineages (Watanabe et al. 2014; Devos et al. 2017; Somerville et al. 2018). To support and extend those findings, our study highlighted the critical role of p63 antagonizing Brn2 in defining more classical squamous cell states within the context of LUSC. Our findings suggest that switching the Sox2 collaborating partner from Brn2 to p63 contributes a shift in cell differentiation states in LUSC via dramatically changing Sox2 cistrome. This phenomenon also supports a role of Sox2 in determining cell differentiation states by having specific partners is maintained in malignant cells. As this switch has not been described in normal differentiation or regeneration of lung cells, whether this transition from Brn2 to p63 is unique to LUSC remains to be investigated. Of note, p63 overexpression in the Brn2-positive ‘neural’ LUSC cells decreased cell growth both *in vitro* and *in vivo*. It suggests that rather than playing a role as a general oncogene, p63 defines more squamous differentiation states and the lineage program of this Brn2-positive ‘neural’ LUSC could lead to more aggressive phenotypes as cancer cells compared to that of the classical LUSC defined by p63. Our study raised a hypothesis that a loss of p63 in the classical LUSC can unveil a latent neural state that is represented by the novel Brn2-high subtype. However, our classical LUSC cell lines were dependent on p63 for their survival and p63 knockout did not induce Brn2 expression in a short period of time. Further studies are needed to explore this hypothesis.

We also observed a change of RTK signaling activity profile through a shift in cell differentiation states induced by p63. It has been reported that p63 regulates a various signaling pathway molecules depending on cell contexts such as β-catenin, EGFR and Jagged1 in human airway epithelial basal cells (Warner et al. 2013), FGFR2 in murine squamous cell carcinoma model (Ramsey et al. 2013) and NRG1 in mammary basal cells (Forster et al. 2014). In our study, we identified several RTKs whose expression and phosphorylation level are altered after p63 overexpression. While functions of positive ErbB4 signaling in the ‘neural’ LUSC cells are unclear, the p63-overexpressed LUSC cells showed more responses to ErbB3/EGFR pathway stimulation and high ERK activity even without stimulation. It is possible that the ‘neural’ LUSC cells are less dependent on AKT and MAPK pathways similarly to SCLC. Future studies are needed to investigate the relationships between lineage programs and networks of signaling pathways and this would help us making therapeutic strategies for each lineage cancer program.

Taken together, we identified a novel ‘neural’ lineage signified by Sox2 and Brn2 in LUSC by investigating its super-enhancer landscape. Brn2 cooperates with Sox2 in determining its transcriptional program, which is overwhelmed and reprogrammed by p63, the classical squamous partner of Sox2. Characterization of each cancer lineage to identify its unique vulnerabilities could lead to a novel approach to make therapeutic strategies for this heterogeneous disease.

## Materials and Methods

### Cell lines

Lung cancer cell lines (HCC95, KNS62, HCC2814, LK2, NCI-H520, SQ-1, EBC-1, HCC2279, NCI-H2887, HCC2450, Calu-1, HARA, LC-1/SQSF) were maintained in RPMI-1640 (Gibco) with 10% FBS and 1% penicillin–streptomycin (Gibco). HEK293T cells were maintained in DMEM with 10% FBS and 1% penicillin–streptomycin (Gibco). Cultured cells were regularly tested for mycoplasma using the mycoAlert Detection Kit (Lonza).

### ChIP-seq

For H3K27ac, Sox2 and p63 ChIP, cells were performed as previously described with modifications (Watanabe et al. 2014). Cells were crosslinked with 1% formaldehyde in PBS for 10 min at room temperature, washed in 5 mg/ml bovine serum albumin (BSA) in PBS and then in just cold PBS, re-suspended in lysis buffer (50 mM Tris-HCl pH 8.1, 10 mM EDTA, 1% SDS, 1× protease inhibitor cocktail (Thermo Fisher Scientific)) and sonicated with the Covaris M220 sonicator or the Diagenode Bioruptor sonicator to obtain chromatin fragment lengths of 100-to-1,000 bp judged by Bioanalyzer DNA High sensitivity kit (Agilent). Fragmented chromatin was diluted in IP buffer (20 mM Tris-HCl pH 8.1, 150 mM NaCl, 2 mM EDTA, 1% Triton X-100) and incubated overnight at 4 °C with Protein G magnetic beads (Dynabeads: ThemoFisher) that had been pre-incubated with anti-H3K27Ac (Abcam, ab4729) or anti-Sox2 (R&D, AF2018) or anti-p63 (Santa cruz, sc-8431) antibodies.

For Brn2 ChIP, cells were crosslinked with 2 mM disuccinimidyl glutarate in PBS with 1 mM MgCl_2_ for 45 min at room temperature and 1% formaldehyde in PBS for 11 min at room temperature, washed in 5 mg/ml BSA in PBS and then in just cold PBS, re-suspended in nuclear extraction buffer (10 mM Tris-HCl pH 7.5, 10 mM NaCl, 3mM MgCl_2_, 0.1 % IGEPAL CA-630, 1× protease inhibitor cocktail) and incubated on ice for 10 min. Extracted nuclei were then washed and resuspended in micrococcal nuclease digestion buffer (0.3 M sucrose, 20 mM Tris-HCl pH 7.5, 3 mM CaCl_2_, 1× protease inhibitor cocktail) and incubated with 12,000 gel units/ml micrococcal nuclease (NEB, M0247S) at 37 °C with frequently mixing to digest chromatin to lengths of approximately 100-to-1,000 bp. Digestion was stopped with 25 mM EDTA pH 8.0 and nuclei was re-suspended in lysis buffer (50 mM Tris-HCl pH 8.0, 10 mM EDTA, 1% SDS, 1× protease inhibitor cocktail) and sonicated with the Diagenode Bioruptor sonicator (low intensity, 30 sec on and 30 sec off, 2 cycles). Fragmented chromatin was diluted in IP buffer (20 mM Tris-HCl pH 8.1, 150 mM NaCl, 2 mM EDTA, 1% Triton X-100) and incubated with anti-Brn2 (Cell Signaling Technology, #12137) antibody overnight at 4 °C. Protein G magnetic beads (Dynabeads: ThemoFisher) were then added and the IP reaction was incubated 2 hours at 4 °C.

Immunoprecipitates were washed six times with wash buffer (50 mM HEPES pH 7.6, 0.5 M LiCl, 1 mM EDTA, 0.7% Na deoxycholate, 1% IGEPAL CA-630) and twice with TE buffer. Immunoprecipitated (or no IP input) DNA was treated with RNase A and Proteinase K on the beads, recovered in 1% SDS and 0.1 M NaHCO3 over a period of 7 h at 65 °C, and purified with DNA clean and concentrator-25 (Zymo Research). Up to 10 ng of DNA was used for the library construction using NEBNext Ultra II DNA Library Prep Kit (NEB, E7645). Sequencing was performed on NextSeq500 (Illumina) for 38 nucleotides each from paired ends according to the manufacturer’s instructions.

### ChIP-seq data analysis

For H3K27ac ChIP, ChIP-enriched regions (peaks) were identified by MACS (Zhang et al. 2008) after aligning to the human reference genome hg19/GRCh37. To define ‘super-enhancers’, we used the ROSE2 pipeline (Lin et al. 2016). H3K27ac enriched peaks were stitched together, using an optimal distance determined by the algorithm per sample up to 12.5 kb. The algorithm generates plots for all stitched enhancers per each sample and defines super-enhancers on the basis of top-ranked enhancers with highest read counts at the tangential cut-off. Unsupervised hierarchical clustering was performed using signals (read counts on defined super-enhancer regions) near transcriptional regulators. A list of the transcriptional regulators was obtained from the genes annotated with gene ontology term, DNA-binding (GO: 0003677). Principle component analysis was performed based on the same signals per region, and top 3 components were depicted in three dimensional plots.

ChIP signals for each sample were visualized on integrative genome viewer (IGV) genome browser (Robinson et al. 2011) using wiggle files with a 10 bp resolution for H3K27ac modification generated by MACS with tag shift that was rescaled to normalize to a total number of uniquely alignable sequences by WigMath function of Java-Genomic Toolkit. Differentially enriched super-enhancers were determined by comparing the ‘neural’ subgroup against the ‘classical’ subgroup using ‘samr’ R package (Tusher et al. 2001), based on cutoffs of fold change>2 and SAM sore>4.

For Sox2, Brn2 and p63 ChIP, peaks were identified by MACS (Zhang et al. 2008) after aligning to the human reference genome hg19/GRCh37. ChIP signals on Sox2, Brn2 and p63 peaks detected on any samples were visualized as a heatmap generated by ‘heatmap’ function in Cistrome analysis pipeline (http://cistrome.org/) or plotHeatmap function of deepTools (Liu et al. 2011) using wiggle files with a 10 bp resolution generated by MACS with tag shift that was rescaled to normalize to a total number of uniquely alignable sequences by WigMath function of Java-Genomic Toolkit.

### RNA-seq

Total RNAs from engineered LK2 cell lines were extracted using the Qiagen RNeasy kit. Poly-adenylated RNA was enriched from 1ug of RNA for each sample with the NEBNext PolyA mRNA Magnetic Isolation Module (NEB, E7490), incubated at 94°C for 15 min and double-strand cDNA was synthesized using SuperScript III reverse transcriptase (ThermoFisher) and NEBNext® Ultra™ II Directional RNA Second Strand Synthesis Module (NEB). Up to 10 ng of cDNA was used for the Illumina sequencing library construction using NEBNext Ultra DNA Library Prep Kit (NEB, E7645). Paired ends sequencing was performed on NextSeq 500 (Illumina) for 38 nucleotides from each end according to the manufacturer’s instructions.

### RNA-seq analyses

RNA-seq gene expression data for *SOX2*, *TP63* and *POU3F2* as well as SNP-array copy number data for *SOX2* were obtained from Cancer Cell Line Encyclopedia (CCLE) (http://www.broadinstitute.org/ccle/home). RNA-seq data of 501 tumor tissues were obtained from TCGA-LUSC dataset (Cancer Genome Atlas Research Network 2012). We downloaded htseq-counts as read counts for each gene in May 2018 (https://portal.gdc.cancer.gov/). The TCGA read counts originally aligned for Ensemble transcripts were converted to corresponding RefGene symbol based on the USCS database. mRNA abundance was estimated from read counts in Transcripts Per Million (TPM) as described in Wagner et el. (Wagner et al. 2012). GTEx TPM matrix (https://gtexportal.org/home/datasets) was downloaded from GTEx data portal in July 2018 (GTEx Consortium 2013). Data from the brain hypothalamus region were used for the analysis. Log_2_-transformed TPM values were used as log_2_(TPM+1) for the following analysis. Two RNA-seq data (EGAD00001001244 and GSE60052) were downloaded for Small Cell Lung Cancer (SCLC) (George et al. 2015; Jiang et al. 2016). We downloaded raw fastq files, aligned to the human reference genome hg19, and estimated mRNA abundance in terms of TPM.

For the TCGA-LUSC RNA-seq data, the 501 LUSC tumor tissues were classified based on the expression level of *TP63*, *SOX2* and *POU3F2*. We used TPM cutoffs using (mean – standard deviation) for *TP63* and *SOX2* as their expressions are normally distributed (Fig. 2C and fig. S2, C and D) and TPM>1 for *POU3F2* since most of the tumors have little expression of the gene (Fig. 2A). Hierarchical clustering of the LUSC tumors (n=416) was performed using the differentially expressed genes between *POU3F2*-high/ *TP63*-low and *POU3F2*-low/ *TP63*-high tumors with complete-linkage clustering (Fig. 2E). Kaplan-Meier curves were plotted to compare survival for patients with *POU3F2*-high*/ TP63*-low tumors versus patients with *POU3F2*-low*/ TP63*-high tumors (Fig. 3C).

To perform RNA-seq analyses for engineered LK2 cell lines, sequencing reads were aligned to the human reference genome hg19/GRCh37 using Tophat (v2.1) and read counts per RefGene symbol on the USCS database was estimated using the htseq-count function in SAMtools (Li et al. 2009). Then DEseq2 (Love et al. 2014) was used to identify the differentially expressed genes between control LK2 cells (two stable GFP-overexpressed LK2 cells and duplicates of inducible DNp63-expressing LK2 cells without doxycycline) and DNp63-overexpressed LK2 cells (two stable DNp63-overexpressed cells and duplicates of inducible DNp63-expressing LK2 cells after treatment with 2 µg/ml doxycycline for 6 days) and to identify genes significantly associated with

To identify potentially enriched functions of selected gene sets of interest, we compared these gene sets with the genes annotated by the same Gene Ontology (GO) terms curated in the Molecular Signature Database (MSigDB) (Subramanian et al. 2005). Each of 5917 GO terms included in “C5” collection (version 6.0 downloaded in 2017) was compared with query gene sets. Any GO terms consisting of more than 2000 genes were considered non-specific and removed from the analysis.

### Immunofluorescence

Cells were grown on glass coverslips coated with poly D-lysine (Neuvitro), fixed in 4% paraformaldehyde for 10 min and permeabilized with 1x PBS containing 0.25% Triton X-100 for 10 min. Primary antibodies were used at the following dilutions: anti-Brn2 at 1:200 (Cell Signaling, #12137), and anti-Sox2 at 1:400 (Abcam, ab171380). Fluorescent signal was detected with secondary antibodies conjugated with Alexa Fluor (ThermoFisher #A-11034, #A-21236) diluted at 1:2000, and coverslips were mounted (ProLong Diamond Antifade Mountant with DAPI: ThermoFisher). Images were obtained with a Leica DM5500 B fluorescence microscope at 20x objective magnification. Images were processed using the Fiji distribution of ImageJ (https://fiji.sc/).

### Western blotting and co-immunoprecipitation

To determine protein expression, cell lysates were prepared by incubating cells in lysis buffer A (150 mM NaCl, 50 mM Tris-HCl at pH 8.0, 1% IGEPAL CA-630, 0.5% Na deoxycholate, 0.1% SDS, protease inhibitors) or lysis buffer B (150 mM NaCl, 40 mM Tris-HCl at pH 8.0, 1% IGEPAL CA-630, 0.5% Na deoxycholate, 2mM EDTA at pH 8.0, protease/phosphatase inhibitors) for 30 min at 4°C. After centrifugation to remove insoluble debris, lysates were immunoblotted with use of an anti-Brn2 antibody (Santa Cruz Biotechnology, sc-393324 or Cell Signaling Technology, #12137), anti-p63 antibody (Santa Cruz Biotechnology, sc-8344, sc-8431 or Cell Signaling Technology, #13109), anti-Sox2 antibody (R&D, AF2018 or Cell Signaling Technology, #3579), anti-ErbB4 antibody (Cell Signaling Technology, #4795), anti-phospho-ErbB4 antibody (Cell Signaling Technology, #4757), anti-ErbB3 antibody (Cell Signaling Technology, #12708), anti-phospho-ErbB3 antibody (Cell Signaling Technology, #4791), anti-EGFR antibody (Millipore, #06-847), anti-phospho-EGFR antibody (Abcam, ab40815) anti-phospho-Akt antibody (Cell Signaling Technology, #4060) and anti-phospho-Erk1/2 antibody (Cell Signaling Technology, #4370), anti-β-Actin antibody (Sigma), or anti-vinculin (Sigma). For western blotting to profile ErbB family signaling, engineered LK2 cells were serum-starved for four hours, and then incubated with anti-ErbB4 monoclonal antibody P6-1 (a kind gift from the Hideyuki Saya laboratory, Keio University) for 15 minutes, followed by stimulation with recombinant human NRG1-β1 EGF domain (R&D) for 30 minutes.

For co-immunoprecipitation, whole cell lysate of cells was prepared in IP lysis buffer (25 mM Tris-HCl pH 7.4, 150 mM NaCl, 1 mM EDTA, 1% IGEPAL CA-630, 5% glycerol, protease inhibitors) followed by incubation with anti-BRN2 antibody, anti-SOX2 antibody, or normal IgG antibody and Dynabeads Protein G (ThemoFisher) on a rotator overnight at 4°C. After washing with PBS + 0.05% Tween 20, the beads were boiled for 10 min in 1x SDS sample buffer. The immunoprecipitates were then processed for immunoblotting.

### Lentiviral introduction of genes

*POU3F2*, *DNp63α* or *GFP* open reading frame (ORF) was cloned into pLEX_306 (a gift from David Root, Addgene #41391), pLEX_307 (a gift from David Root, Addgene #41392) or pLIX_403 (a gift from David Root, Addgene #41395) using Gateway® cloning methods according to manufacturer’s recommendations. For lentiviral vectors production, HEK293T cells were seeded in 10-cm tissue culture dish and incubated at 37°C and 5% CO_2_. Cells at 80% confluency were co-transfected with 10 µg of lentiviral expression constructs, 7.5 µg of psPAX2 and 2.5 µg pMD2.G vectors using TransIT-Lenti (Mirus) following manufacturer’s recommendations. At 48 h post transfection, supernatants were collected, filtered (0.45 µm) and stored at −80°C. Cells were infected with supernatant containing lentivirus supplemented with polybrene at a final concentration of 8 µg/mL and then selected with puromycin (2-3 µg/mL for 4-6 days). Ectopic protein expression was confirmed via immunoblotting.

### CRISPR-Cas9 genome editing

Cells stably expressing human codon-optimized S. pyogenes Cas9 were generated by infection with the lentiCas9-Blast plasmid (a gift from Feng Zhang, Addgene, # 52962). sgRNAs were cloned at BbsI site downstream of the human U6 promoter in a lentiviral vector containing eGFP downstream of the human PGK promoter (a kind gift from the Brian Brown laboratory, Icahn School of Medicine at Mount Sinai). Lentivirus was produced as above. Cells were first infected with the lentiCas9-Blast lentivirus, and then selected with blasticidin (5 µg/mL for 10 days) for cells expressing the Cas9 nuclease. Cells were then infected with pLenti-GFP-sgRNA. sgRNA target sequences were selected from Brunello library(Doench et al. 2016) and Orzol et al. (Orzol et al. 2016). Non-target sgRNAs from the Gecko library v2 (Sanjana et al. 2014) were used as scramble sgRNAs. sgRNA target sequences are listed in Supplemental Table S4.

### Immunohistochemistry

Immunohistochemical analyses were performed for xenograft tumor specimens and human primary LUSC tumor specimens. Total 10 FFPE specimens of human primary LUSCs resected from patients at Icahn School of Medicine at Mount Sinai (ISMMS, USA) were obtained and cut into 5-µm-thick sections.

Immunohistochemical staining for Brn2, after being deparaffinized and rehydrated, FFPE tissue slides (5 µm thick) were heated at 95°C in 10 mM citrate buffer (pH 6.0) for antigen retrieval for 30 minutes. The sections were incubated with 0.3% H_2_O_2_ in TBS for 15 minutes to block endogenous peroxidase activity and were incubated with 10% normal goat serum (Jackson ImmunoResearch) in TBS for 30 minutes to block non-specific staining. After rinsing with TBS + 0.025% Triton X-100, the sections were incubated with anti-Brn2 antibody (1:100; Cell Signaling #12137) at 4°C overnight, followed by incubation with biotin-conjugated Goat anti-Rabbit secondary antibody (Vector Laboratories) at room temperature for 1 hour. Then, the sections were incubated with the ABC reagent (Vector Laboratories) and visualized with ImmPACT-DAB Peroxidase Substrate (Vector Laboratories). Immunohistochemical staining for Sox2 and p63 was conducted using a Leica Bond-III automated slide stainer. The sections were deparaffinized and subjected to heat induced antigen retrieval with high pH EDTA buffer for 20 minutes and 10 minutes, respectively. The sections were incubated with anti-Sox2 antibody (prediluted; Cellmarque clone SP76) or anti-p63 antibody (prediluted; Biocare Medical, clone 4A4) as per manufacturer’s directions. All slides were counterstained with hematoxylin before being mounting.

### Cell proliferation assay

Cells were plated onto sextuplicate wells of a 96-well plate; three identical plates were prepared. Cell proliferation was assayed at 24, 72 and 120 h after plating with alamarBlue Cell Viability Reagent (ThermoFisher) and fluorescence at 585 nm was measured on a Spectra Max3 plate reader (Molecular Device) according to the manufacturer’s protocol at excitation of 555 nm. Cell viability at 72 and 120 h were corrected for the ratio to control cells from the 24-h reading to account for plating unevenness.

### Cell cycle analysis

Cells were fixed with 70 % ethanol at least overnight at 4°C. After washing with PBS, cells were incubated in PBS containing 100 µg/ml RNase and 50 µg/ml propidium iodide overnight at 4°C overnight. DNA content was analyzed by FACS Canto II (BD Bioscience), and quantitative analyses for the proportions of cells in cell cycle were performed using FlowJo software.

### Xenograft model

Engineered LK2 cells (4 × 10^6^ cells) were injected with a 1:1 mixture of 50 µl cell suspension and 50 µl Matrigel (Corning) subcutaneously into both flank regions of 4- to 6-week-old male NOD-scid or NOD-scid gamma mice (Jackson Laboratory). Tumor volume (length × width^2^ /2) was measured twice a week. Before tumor size reached at 1000 mm^3^, mice were sacrificed and tumors were immersed in formalin for immunohistological analysis. All animal procedures and studies were approved by the Mount Sinai Institutional Animal Care and Use Committee (IACUC) (protocol number, IACUC-2018-0021).

### Phospho-RTK array analysis

The Human Phospho-RTK Array Kit (R&D Systems) was used to determine the relative levels of tyrosine phosphorylation of 42 distinct RTKs, according to the manufacturer’s protocol. 300 µg of total protein was used for each membrane. Chemiluminescent signals were captured with a Chemidoc MP Imaging System (Bio-Rad laboratories) and images were processed using Image Lab software (Bio-Rad laboratories).

### Statistical analyses

Differentially enriched super-enhancers were determined using ‘samr’ R package (Tusher et al. 2001), based on cutoffs of fold change>2 and SAM sore>4. For the TCGA-LUSC RNA-seq data, Pearson’s correlation coefficient between gene expression levels was calculated and associated *P*-value was determined. *T*-test was performed to identify differentially expressed genes between *POU3F2*-high/ *TP63*-low and *POU3F2*-low/ *TP63*-high tumors based on cutoffs of fold change>2 and FDR<0.01. Significance of the overlap between gene sets was determined using Fisher’s exact test. To compare the overall survival, Kaplan-Meier curves were plotted and log-rank test was performed. DEseq2 (Love et al. 2014) was used to identify the differentially expressed genes between control and DNp63-overexpressed LK2 cells based on cutoffs of fold change>1.5 and FDR<0.05. In addition, likelihood ratio test model was used to identify genes significantly associated with conditions based on cutoffs of fold change>2 and FDR<0.01. For cell proliferation assay, xenograft tumor growth measurement and cell cycle analysis, differences between two groups were examined using two-tailed *t*-test. Bonferroni corrections were performed in case multiple comparisons were conducted.

## Acknowledgements

We thank Chiara Vardabasso, Dan Hasson, Katsutoshi Sato, Aleksandra Wroblewska, Almuneda Bosch and Iman Tavassoly for helpful discussions; Aleksandra Wroblewska and Brian D. Brown for providing a lentiviral vector for cloning sgRNAs; NextSeq Sequencing Facility of the Department of Oncological Sciences at ISMMS, Saboor Hekmaty, Irene Salib, Gayatri Panda and Ravi Sachidanandam for sequencing assistance; and Stephanie Tuminello and Michael William for technical assistance. The authors also thank the Flow Cytometry Core facility and the Biorepository and Pathology Core Facility at ISMMS. This work was supported in part through Tisch Cancer Institute at ISMMS and the computational resources and staff expertise provided by Scientific Computing at ISMMS. This study was supported by grants from the 2017 ATS Foundation Unrestricted Grant: Pulmonary (H. Watanabe) and the American Lung Association of the Northeast Lung Cancer Discovery Award (H. Watanabe). T. Sato is supported by the Japanese Respiratory Society the 6th Lilly Fellowship Program and the Uehara Memorial Foundation. R. Kong is supported by Shaanxi Provincial Natural Science Foundation, China (No. 2017JM8046).

## Author contributions

T.S. and H.W. conceived and designed the study. T.S., S.Y., R.K., M.F., A.S., P.C., A.P., and M.B.B. conducted experiments and/or acquired data. T.S., S.Y., M.B.B., and H.W. analyzed data. O.N., T.M., M.B.B., and C.A.P. provided administrative, technical and/or material support. C.A.P., J.Z., and H.W. supervised the study. T.S., S.Y., and H.W. wrote, reviewed, and revised the manuscript. All authors reviewed and commented on the manuscript.

