## Supplementary Information for "Epigenomic profiling discovers trans-lineage SOX2 partnerships driving tumor heterogeneity in lung squamous cell carcinoma"

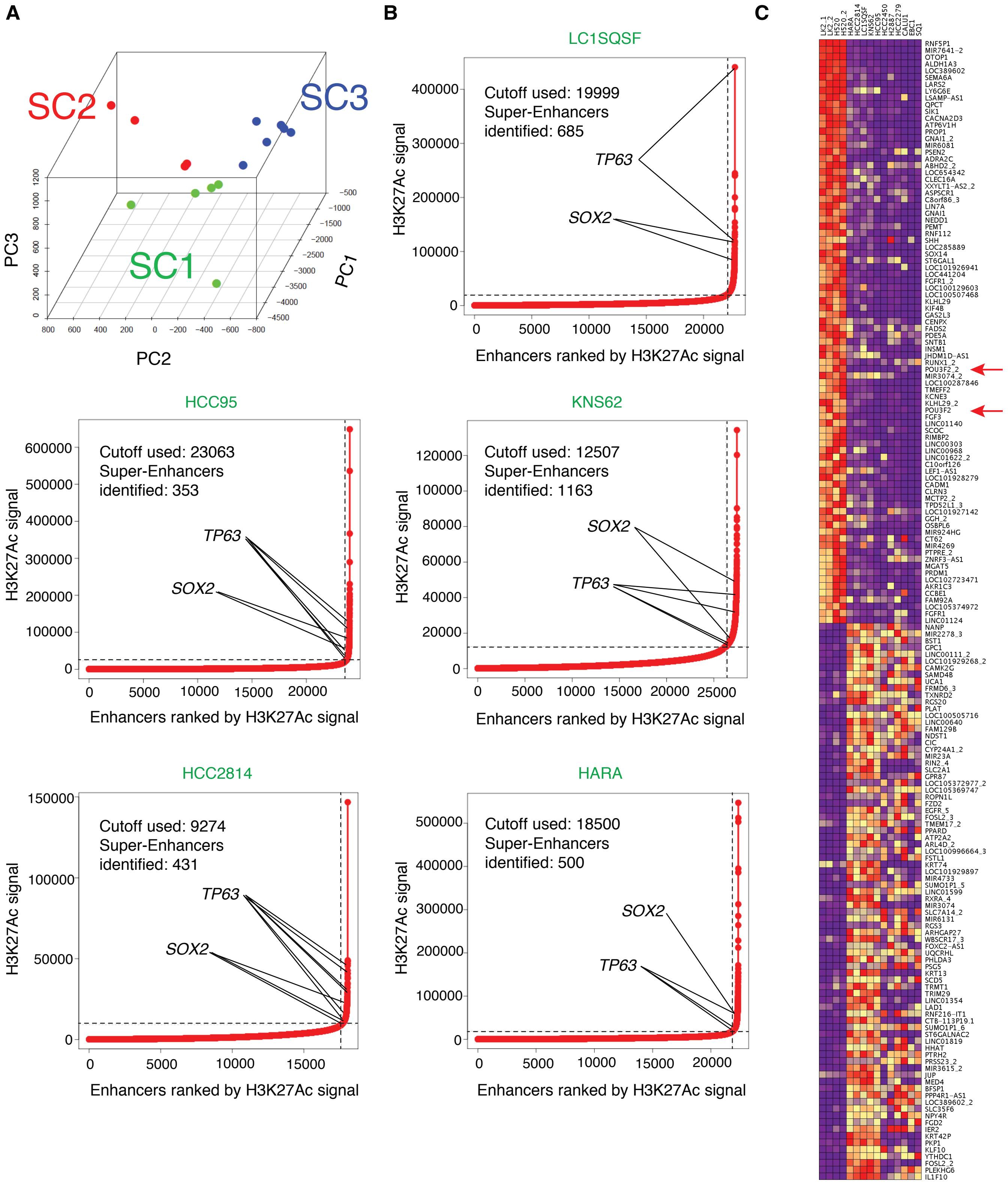


**Supplemental Figure S1.** Super-enhancer profiling identifies three subtypes of LUSC. *(A)* Principal component analysis of super-enhancer profiles near transcriptional regulator genes in 13 LUSC cell lines. Each point represents a LUSC cell line that is colored based on unsupervised hierarchical clustering using super-enhancer scores near transcriptional regulator genes (Fig. 1A). *(B)* Super-enhancer plots using H3k27ac scores in the classical subgroup of LUSC cell lines. *(C)* Super-enhancer loci with differential super-enhancer scores near transcriptional regulator genes in the small subset of LUSC cell lines compared to the classical subgroup of LUSC cell lines.

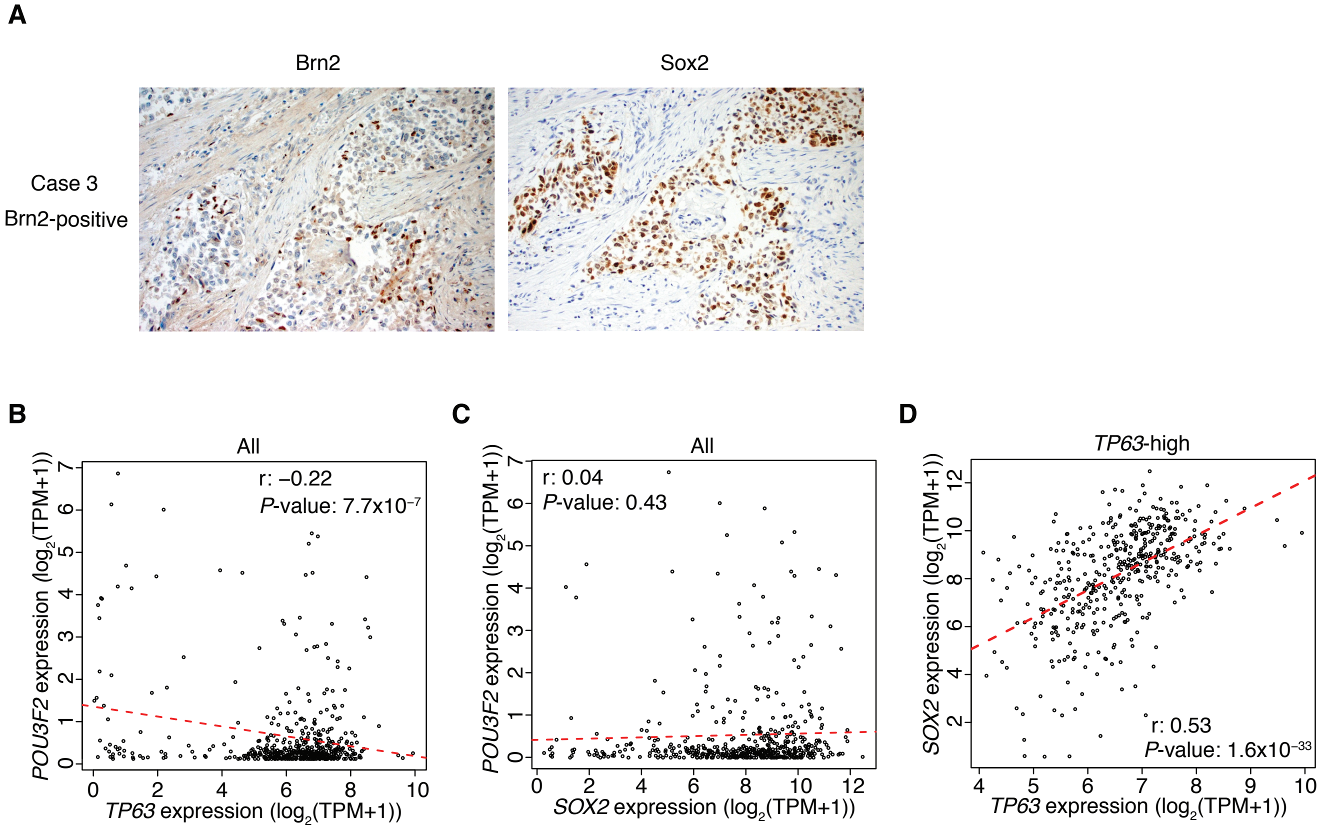


**Supplemental Figure S2.** Brn2 is expressed in a subset of human primary LUSC tumors. *(A)* Immunohistochemical staining of Brn2 and Sox2 and H&E staining in Brn2-positive and negative human LUSC tumors. Representative images are shown (original Images, ×200). *(B)* Scatter plots of expression of *TP63* and *POU3F2* among the entire dataset of TCGA-LUSC tumors. *(C)* Scatter plots of expression of *SOX2* and *POU3F2* among the entire dataset of TCGA-LUSC tumors. *(D)* Scatter plots of expression of *TP63* and *SOX2* in TP63-high LUSC tumors from TCGA.

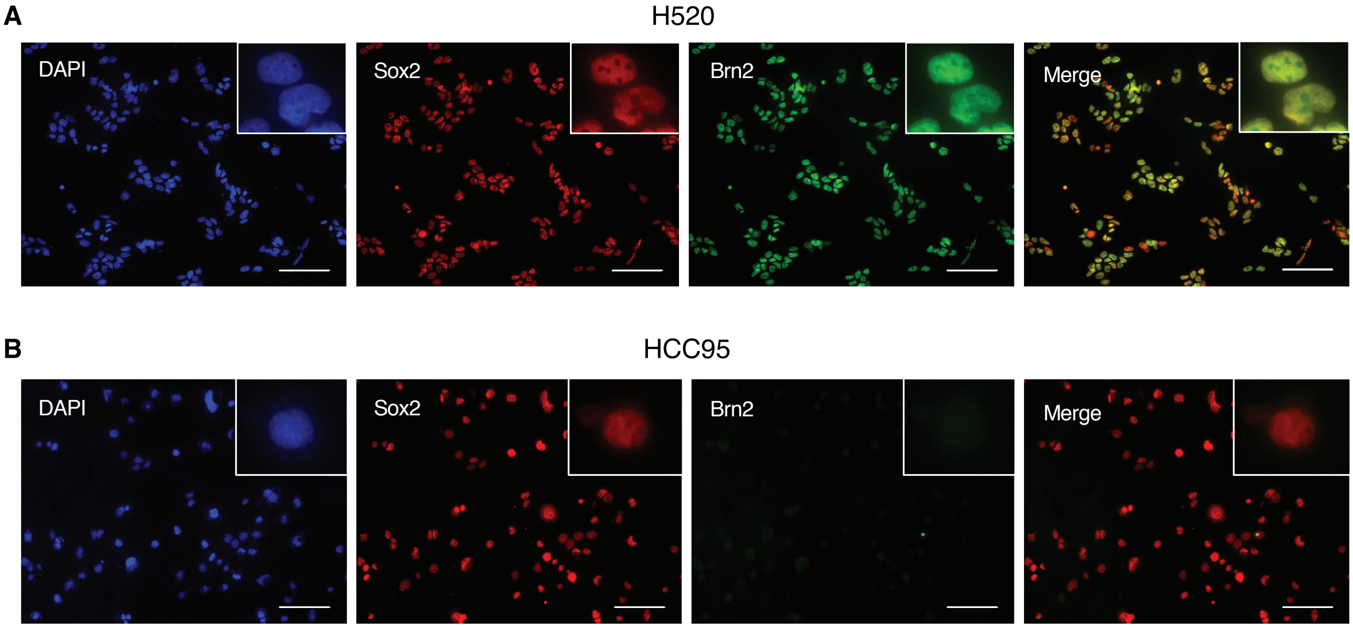


**Supplemental Figure S3.** Brn2 and Sox2 co-localize in nuclei of the 'neural' LUSC cells. *(A,B)* Expression of endogenous Brn2 and Sox2 in NCI-H520 cells *(A)* and HCC95 cells *(B)*, determined by immunofluorescence with anti-Sox2 (*green*) and anti-Brn2 (*red*) antibodies, respectively. DAPI staining (nuclei; *blue*) and merged images are also shown. Original magnification, ×200. Scale bar, 100 µm.

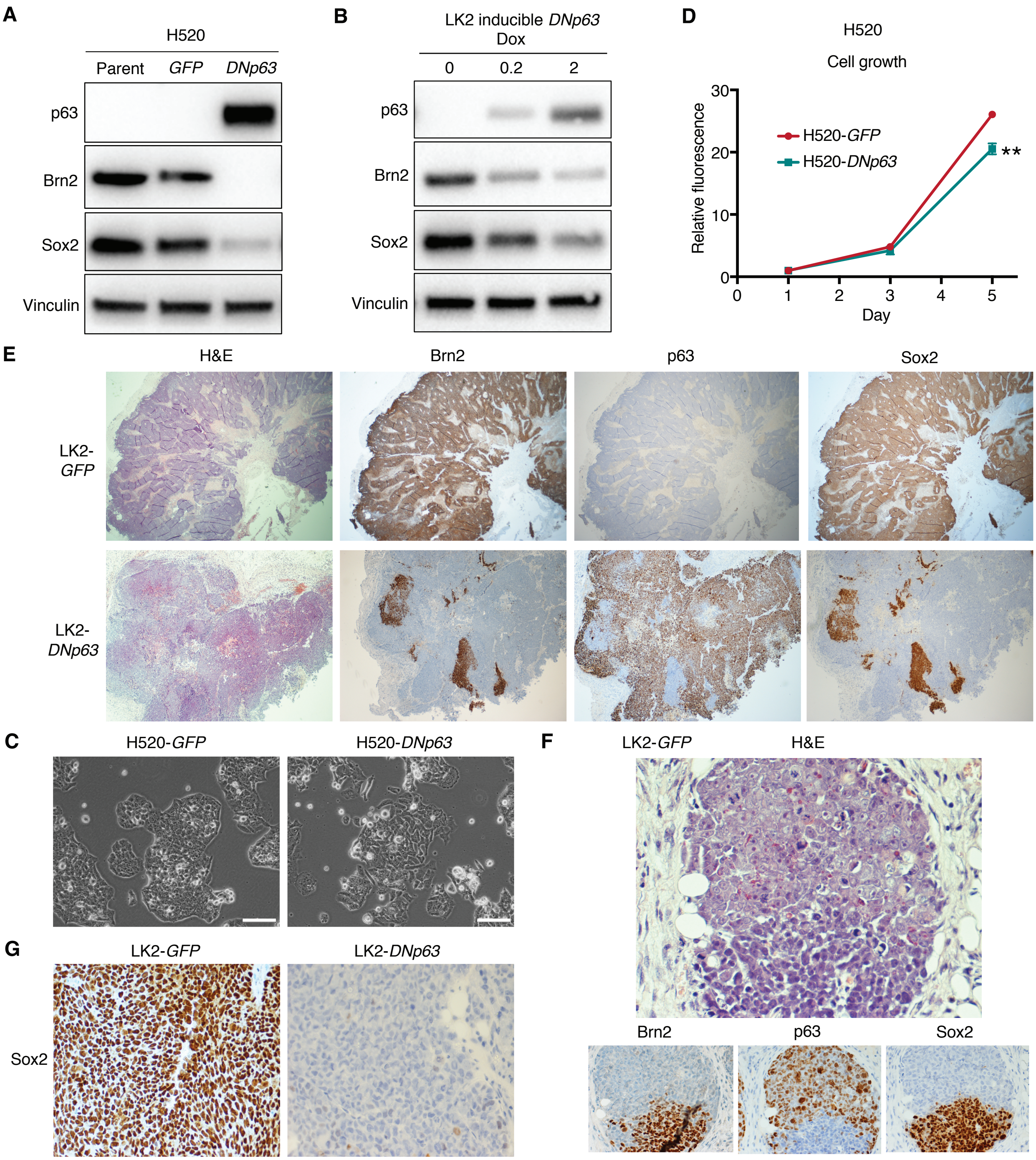


**Supplemental Figure S4.** DNp63 overexpression in the 'neural' LUSC cells suppresses Brn2 expression and induces phenotypic changes. *(A)* Protein expression of p63, Brn2, Sox2 and vinculin as a loading control in parental, GFP-overexpressed and DNp63-overexpressed NCI-H520 cells. *(B)* Protein expression of p63, Brn2, Sox2 and vinculin as a loading control in doxycycline-inducible DNp63-overexpressing LK2 cells. Cells were treated with doxycycline (Dox) for 11 days at the indicated concentrations. *(C)* Phase-contrast microphotographs of GFP-overexpressed and DNp63-overexpressed NCI-H520 cells. Bar = 100 µm. *(D)* Cell growth of GFP-overexpressed and DNp63-overexpressed NCI-H520 cells. Mean ± SD of sextuplicates are shown. **, $P$<0.001 vs. GFP-overexpressed NCI-H520 cells, $t$-test. *(E)* H&E staining and immunohistochemical staining of Brn2, p63 and Sox2 in GFP-overexpressed or DNp63-overexpressed LK2 xenograft. Original Images, ×40. *(F)* H&E staining and immunohistochemical staining of Brn2, p63 and Sox2 in the areas where both p63-positive and negative tumor cells were found in the DNp63-overexpressed LK2 xenograft. Original Images, ×400. *(G)* Immunohistochemical staining of Sox2 in the DNp63-overexpressed LK2 xenograft. Original Images, ×400.

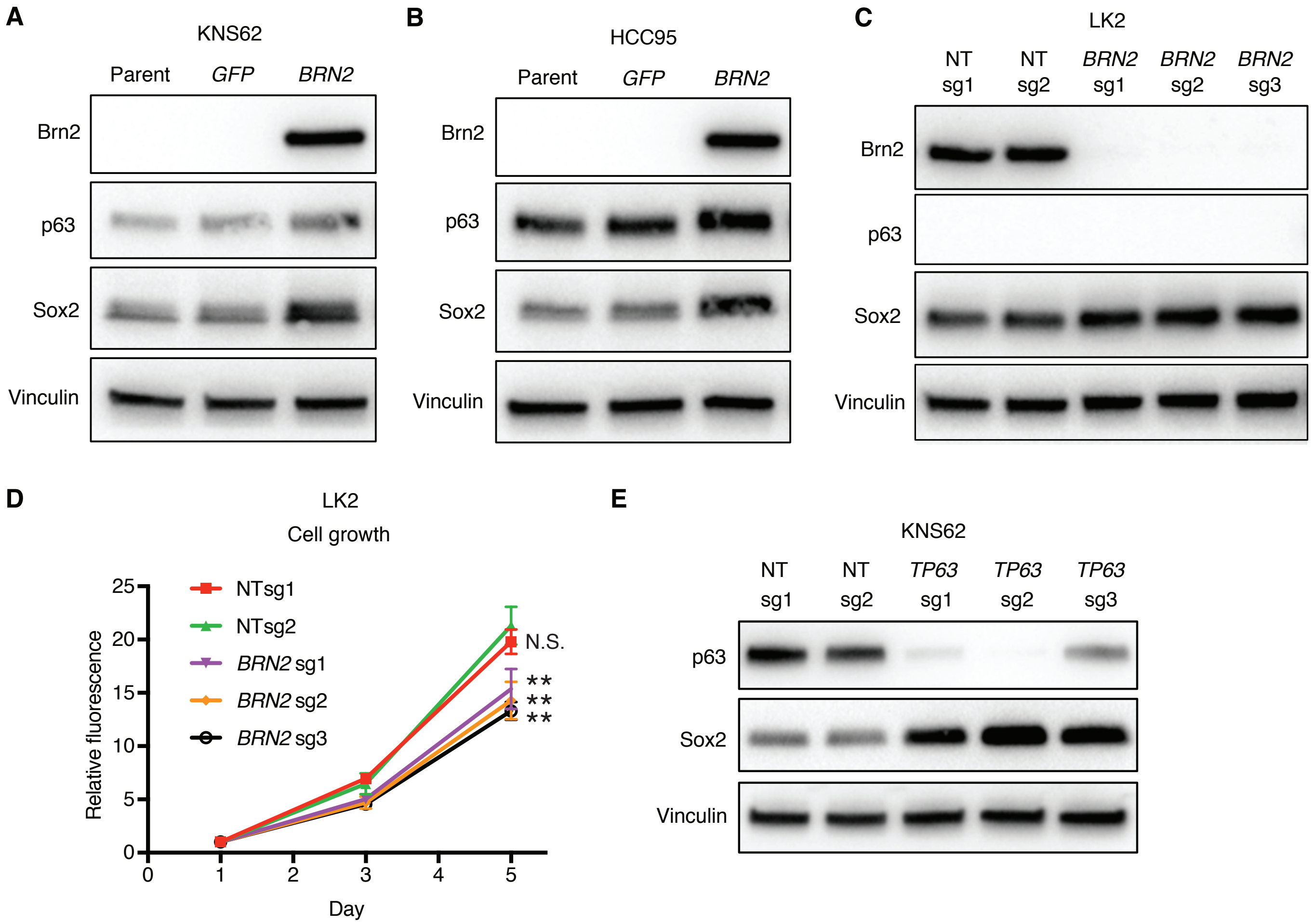


**Supplemental Figure S5.** Brn2 overexpression and p63 ablation in the 'classical' LUSC cells and Brn2 ablation in the 'neural' LUSC cells. *(A,B)* Protein expression of Brn2, p63, Sox2 and vinculin as a loading control in parental, GFP-overexpressed and DNp63-overexpressed KNS62 cells *(A)* or HCC95 cells *(B)*. *(C)* Protein expression of Brn2, p63, Sox2 and vinculin in LK2 cells infected with 2 independent non-target sgRNAs (NT sg1 or 2) or 3 independent *BRN2* sgRNAs (BRN2 sg1, 2 or 3). *(D)* Cell growth of LK2 cells infected with each non-target sgRNA or *BRN2* sgRNA. Mean ± SD of sextuplicates are shown. **, $P<0.001$ and N.S., $P>0.05$ vs. LK2 cells with non-target sgRNA2, *t*-test with Bonferroni correction. *(E)* Protein expression of p63, Sox2 and vinculin in KNS62 cells infected with 2 independent non-target sgRNAs (NT sg1 or 2) or 3 independent *TP63* sgRNAs (*TP63* sg1, 2 or 3).

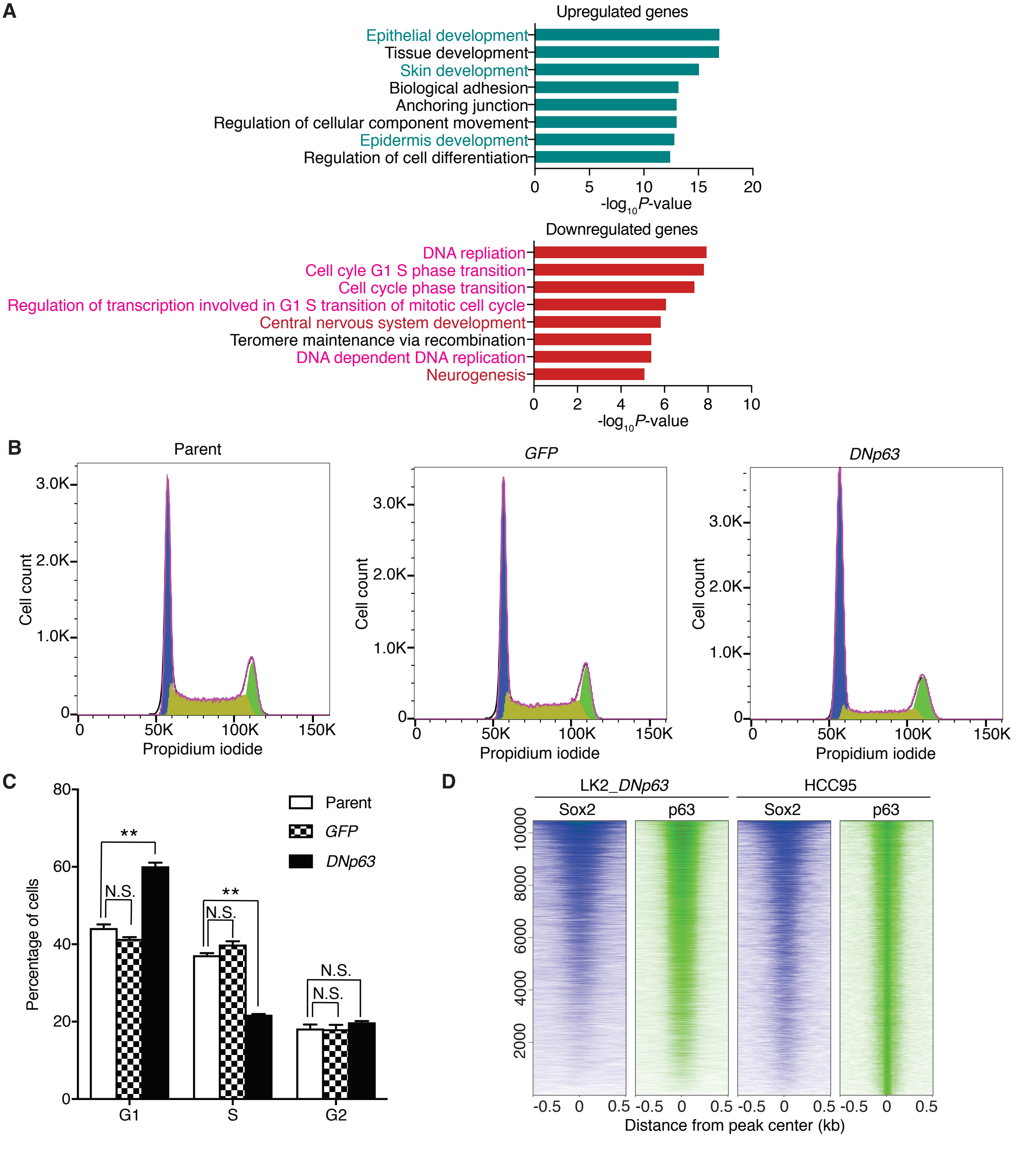


**Supplemental Figure S6.** DNp63 induces a classical squamous-cell transcriptional program and G1 phase arrest in the 'neural' LK2 cells. *(A)* Gene ontology analyses for the differentially up-regulated (*top*) and down-regulated (*bottom*) genes upon DNp63 overexpression in LK2 cells. Likelihood ratio test model was used to identify genes significantly associated with conditions (control -> inducible overexpression -> stable overexpression) based on cutoffs of fold change>2 and FDR<0.01. Enriched functions for these genes are identified based on Fisher's exact test against GO terms curated in MSigDB. *(B)* Propidium iodide staining followed by flow cytometry was used to analyze cell cycle distribution in parental (*left*), GFP-overexpressed (*middle*) and DNp63-overexpressed (*right*) LK2 cells. *(C)* The percentages of cells in G1, S, and G2 phase of cell cycle were measured using flow cytometry after propidium iodide staining in parental, GFP-overexpressed and DNp63-overexpressed LK2 cells. Each bar represents the mean ± SD of triplicate measurements. **, $P<0.001$ and N.S., not significant ($P>0.05$), *t*-test with Bonferroni correction. *(D)* Heatmap depicting analysis of ChIP-seq signals for Sox2 and p63 in DNp63-overexpressed LK2 cells and HCC95 cells at all p63 peak loci. ChIP-seq signal intensity is shown by color shading.

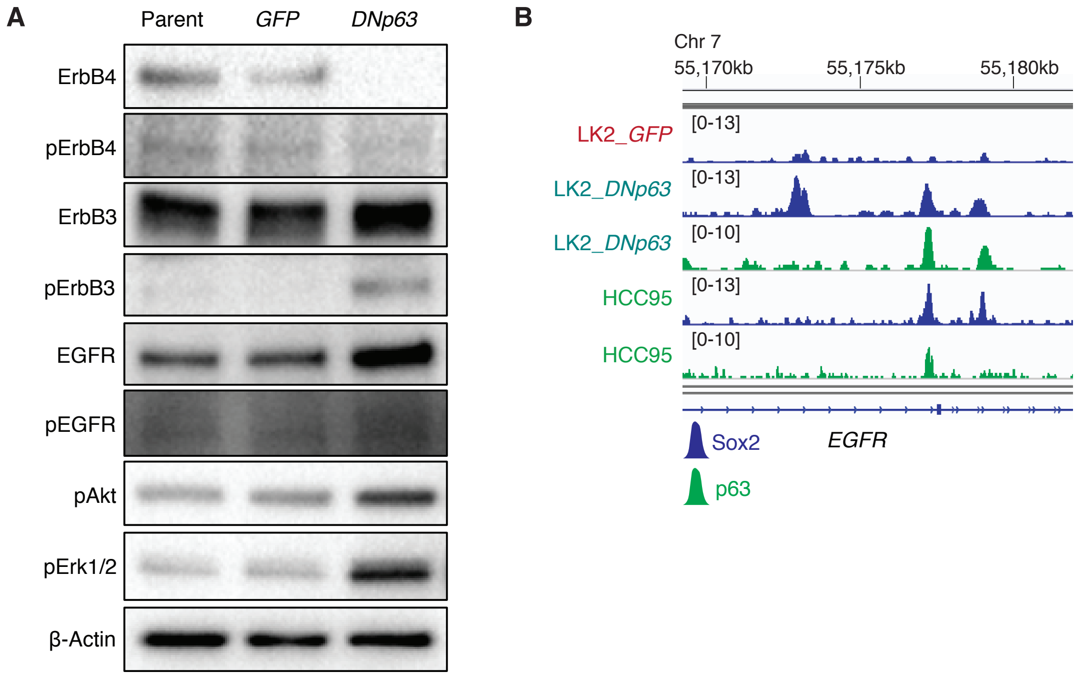


**Supplemental Figure S7.** DNp63 alters ErbB family signaling in the 'neural' LK2 cells. *(A)* Protein expression of ErbB4, phospho-ErbB4, ErbB3, phospho-rbB3, EGFR, phospho-EGFR, phospho-Akt and phospho-Erk1/2 and β-Actin as a loading control in Parental, GFP-overexpressed and DNp63-overexpressed LK2 cells. Cells were cultured with 10% FBS. *(B)* Genome view tracks of ChIP-seq signals for Sox2 and p63 at *EGFR* locus in GFP-overexpressed and DNp63-overexpressed LK2 cells and HCC95 cells.

**Supplemental Table S1.** Gene loci for super-enhancers near transcriptional regulators shared by all the LUSC cell lines in 'classical' subgroup.

| Gene |
|---|
| *SOX2* |
| *TP63* |
| *HES1* |
| *MYC* |
| *NFE2L2* |
| *POU5F1B* |
| *TBL1XR1* |
| *TGIF1* |

**Supplemental Table S2.** Gene loci for super-enhancers with highest signals near transcriptional regulators in LK2 and NCI-H520 cells.

| Rank | LK2 | NCI-H520 |
|---|---|---|
| 1 | ***SOX2*** | *BRD7* |
| 2 | *HOXA10* | *NR2E3* |
| 3 | ***POU3F2*** | *TCF4* |
| 4 | *DLX5* | ***SOX2*** |
| 5 | ***SOX2*** | *ZNF217* |
| 6 | *TERF1* | *ETV5* |
| 7 | *PROP1* | *RCAN1* |
| 8 | *TNRC18* | *NR2E3* |
| 9 | *ARID2* | *TBL1XR1* |
| 10 | *TBX3* | *SOX2* |
| 11 | *HIST2H2AA4* | ***POU3F2*** |
| 12 | *TGIF1* | *TCF4* |
| 13 | *TBL1XR1* | *SATB2* |
| 14 | *RUNX2* | *DBX1* |
| 15 | *WNT5A* | *LHFP* |
| 16 | *TBX2* | *KLF4* |
| 17 | *RUNX1* | *ETV5* |
| 18 | *TRPS1* | *BARX2* |
| 19 | *ZFHX3* | ***POU3F2*** |
| 20 | *KLF4* | *RORA* |

**Supplemental Table S3.** Associations between gene signatures from DNp63-overexpressed LK2 model and those from human TCGA-LUSC dataset based on *POU3F2*/ *TP63* expression.

| Number of genes overlapped/ Odds ratio/ *P*-value | TCGA *POU3F2*-low/ *TP63*-high | TCGA *POU3F2*-high/ *TP63*-low |
|---|---|---|
| Up-regulated genes in DNp63-overexpressed LK2 | 7/ 1.91/ 0.106 | 74/ 9.81/ $2.07 \times 10^{-41}$ |
| Down-regulated genes in DNp63-overexpressed LK2 | 13/ 6.17/ $7.32 \times 10^{-7}$ | 7/ 1.06/ 0.842 |

**Supplemental Table S4.** sgRNA sequences for the CRISPR/Cas9 system.

| | sgRNA sequence (5’-3’) | PAM sequence |
|---|---|---|
| *POU3F2*_sg1 | ATCACCGCGCTGTCCCACGG | CGG |
| *POU3F2*_sg2 | CAAACTGGGATTTACCCAAG | CGG |
| *POU3F2*_sg3 | CACGCCGCTAACCACCACCC | GGG |
| *TP63*_sg1 | CCGTGACGCTGTTCTGCGCG | TGG |
| *TP63*_sg2 | CAATGATTAAAATTGGACGG | CGG |
| *TP63*_sg3 | GCAGTCGAGCACCGCCAAGT | CGG |
| non-target sg1 | ACGGAGGCTAAGCGTCGCAA | N/A |
| non-target sg2 | CGCTTCCGCGGCCCGTTCAA | N/A |
